# An ultra-long acting insulin enables glucose-synchronised release

**DOI:** 10.1101/2025.10.12.681846

**Authors:** Yang Zhang, Juan Zhang, Kangfan Ji, Shaoqian Mei, Chen Su, Yanfang Wang, Guangzheng Xu, Hui Zhang, Junhuan Lin, Xuehui Huang, Xiuwen Zhang, Kairui Xu, Jianchang Xu, Leihao Lu, Mowei Zhou, Wei Huang, Jian Yao, Chuhuan Jiang, Junjie Yan, Xiangsheng Liu, Peifeng Liu, John B Buse, Shiming Zhang, Jinqiang Wang, Zhen Gu

## Abstract

Delayed and weak glucose-responsive kinetics, coupled with short plasma exposure and complicated formulation, remain major obstacles to the clinical translation of glucose-responsive insulin formulations. Here we report a rapidly absorbable, ultra-long acting, and glucose-responsive insulin analogue *via* engineering recombinant human insulin with two phenylboronic acids. The insulin analogue has a long circulating half-life (t_1/2*β*_ ∼150 h) due to its unique, unprecedented, glucose-responsive binding to blood circulation-associated proteins. This interaction drives the formation of a unique systemic reservoir that allows synchronised dynamic response in insulin levels to glucose fluctuations. In type 1 diabetic mice and minipigs, a single subcutaneous dose can maintain normoglycaemia for over one week (47.5-fold compared to insulin glargine). After the last injection of four consecutive weekly administrations in mice, the glucose-lowering effect can last 800 hours. Administering this glucose-responsive insulin analogue with a fully automated insulin delivery system can leverage dual closed loops and achieve a glucose time in range of over 85% and a coefficient of variation of approximately 30%, better than the recommended targets for human therapy. Importantly, no toxicity was identified in the study.

## Introduction

For people with type 1 diabetes and those with advanced type 2 diabetes, insulin replacement therapy is an essential and effective strategy for regulating blood glucose (BG) within the target range ^1,2^. The narrow therapeutic index of insulin, coupled with dynamic insulin needs, requires precise and real-time insulin level adaptation ^3^. As a result, multiple pre-meal insulin injections are used in combination with daily basal insulin to mimic the natural rhythm of insulin secretion. Recently, the introduction of insulin icodec to the European Union market has brought diabetes care into the era of weekly formulations of basal insulin ^4^. Despite these efforts, hypoglycaemia remains a common and sometimes fatal complication. Fear of insulin-induced hypoglycaemia leads to reduced insulin doses, often resulting in suboptimal glucose control and a high risk of long-term hyperglycaemia-related complications ^5^.

Hypoglycaemia primarily stems from either a BG-irrelevant insulin release rate or BG-independent insulin activity. Contrary to conventional insulin, glucose-responsive insulin (GRI) can respond to glycaemic fluctuations and adjust its activity or concentration accordingly. Two major types of GRI, carrier-based ^6–21^ and modification-based GRI ^22–29^, have been developed since the 1970s, and have achieved significant progress recently ^29^. These GRI have demonstrated their glucose-responsive performance mainly *in vitro* and partially in animal studies *via* glucose tolerance tests or clamp studies. However, short circulatory half-lives, inadequate glucose-responsive kinetics, biocompatibility concerns regarding carriers, and delayed time to peak insulin concentrations in response to glucose excursions have not been fully addressed ^30,31^. These issues have been faced in animal studies or clinical trials, and thus no GRI has been clinically approved ^32^. Addressing these issues with current approaches and knowledge remains challenging.

In this study, we have developed a GRI analogue, designated insulin-129, by covalently conjugating 4-carboxy-3-fluorophenylboronic acid (FPBA) to recombinant human insulin (RHI) at A1 and B29 sites (**Fig. 1a**). Insulin-129 exhibits rapid absorption and engages in a novel and unprecedented dynamic binding with immunoglobulin G1 (IgG1) and nostrin, rather than the traditional glycosylated proteins or albumin (**Fig. 1b**). This unprecedented interaction extends circulatory half-life (t_1/2*β*_ ∼150 h), promotes vascular adhesion, and most importantly is glucose-modulable, and therefore facilitates the creation of a glucose-responsive reservoir in systemic circulation (**Fig. 1b**). In streptozotocin (STZ)-induced type 1 diabetic mice, a single subcutaneous (*s.c.*) administration of insulin-129 maintains normoglycaemia for over one week, with no incidence of hypoglycaemia. Upon glucose injection, a more than 3-fold synchronised increase in insulin level is observed, which is associated with a 2-fold temporal change in blood glucose clearance determined by clamp studies. Similarly, in type 1 diabetic minipigs, one injection sustains BG within the normal range for more than a week, with minimal risk of hypoglycaemia. Notably, synchronised fluctuations in insulin-129 levels and blood glucose concentrations are confirmed in minipigs. Capitalising on these unique attributes, insulin-129 is administered with a continuous glucose monitoring system (CGMS), a bespoke but simplified algorithm, and an insulin pump to construct a fully automated insulin delivery system (AID). When evaluated in type 1 diabetic minipigs without human intervention, the AID extends glucose time in range, stabilises glycaemic control and prevents hypoglycaemia. Of note, no significant toxicity has been observed, underscoring its potential safety and efficacy. This platform can be extended to other proteins to prepare long-circulating and glucose-responsive formulations.

**Fig. 1.**
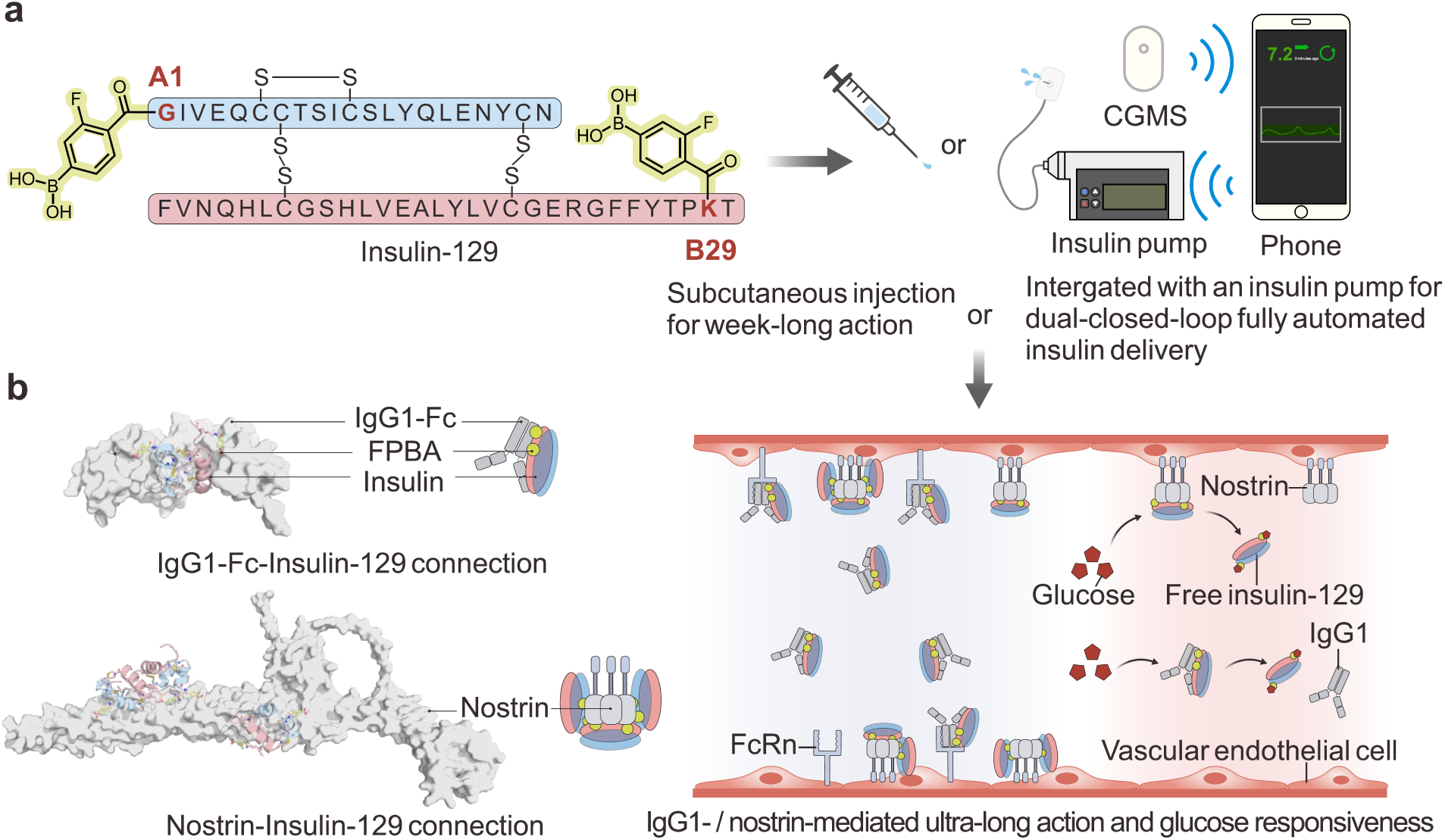
| Chemical structure and glucose-responsive mechanism of insulin-129. **a,** Insulin-129 with dual conjugation of the 4-carboxy-3-fluorophenylboronic acid at the A1 and B29 sites for subcutaneous injection or integration with an insulin pump to achieve dual-closed-loop treatment. CGMS, continuous glucose monitoring system. **b,** The predicted binding configurations of insulin-129 with immunoglobulin heavy constant gamma 1 (IgG1-Fc) and nostrin (left). Immunoglobulin G1 and nostrin mainly mediate the ultra-long action and glucose responsiveness of insulin-129 (right).

### Preparation and characterization of insulin-129, insulin-1, and insulin-29

We mixed FPBA and RHI in the presence of coupling agents and obtained insulin-129 after purification with preparative liquid chromatography (**Fig. 1a and Supplementary Fig. 1a**). Insulin-129 had a m/z value of 6134.72 as determined by high-resolution mass spectrometry (**Supplementary Fig. 1b,c**), indicating two FPBA residues conjugated to one RHI. Tandem mass spectrometry (MS/MS) analysis further confirmed the FPBA modification sites at A1-glycine and B29-lysine (**Supplementary Fig. 1d,e**). The dual conjugation of FPBA did not affect the insulin’s secondary structure as characterized by circular dichroism (**Supplementary Fig. 1f**). After freeze-drying, insulin-129 exhibited a loose, porous structure (**Supplementary Fig. 1g,h**). We also prepared insulin-1 and insulin-29 with FPBA conjugation only at the A1 or B29 sites, respectively (**Supplementary Fig. 1i,j**). Their modifications were likewise verified through MS/MS analysis (**Supplementary Fig. 2k,l**).

### *In vitro* evaluation of glucose binding and discovery of insulin-129’s prolonged duration

Native mass analysis was used to assess the *in vitro* glucose-binding properties of insulin-129, insulin-1, and insulin-29. Insulin-129, possessing two glucose-binding sites, displayed a dissociation constant (*K*_D_) of 6.40 mM (**Fig. 2a**)—well within the normal blood glucose range and similar to that reported in literature ^33^. As a comparison, insulin-1 and insulin-29 bound to glucose with *K*_D_ values of 13.4 mM and 16.4 mM, respectively (**Fig. 2a**). RHI only displayed nonspecific binding to glucose at high glucose concentrations (**Fig. 2a**). Glucose-binding kinetics of insulin-129 in the presence of 10 mM glucose revealed a binding half-life of just 1.60 minutes (**Fig. 2b**), underscoring its potential for a rapid response to glucose fluctuations. Insulin-129 exhibited a similar activity to RHI in activating the insulin receptor in the HepG2 cell line (**Supplementary Fig. 2a,b**). The preliminary treatment efficacy in STZ-induced type 1 diabetic mice validated the longest duration of insulin-129 among the three insulin analogues (**Supplementary Fig. 2c**). Given the glucose binding capability, insulin receptor activating capability, and treatment duration, insulin-129 was selected in the following studies.

**Fig. 2.**
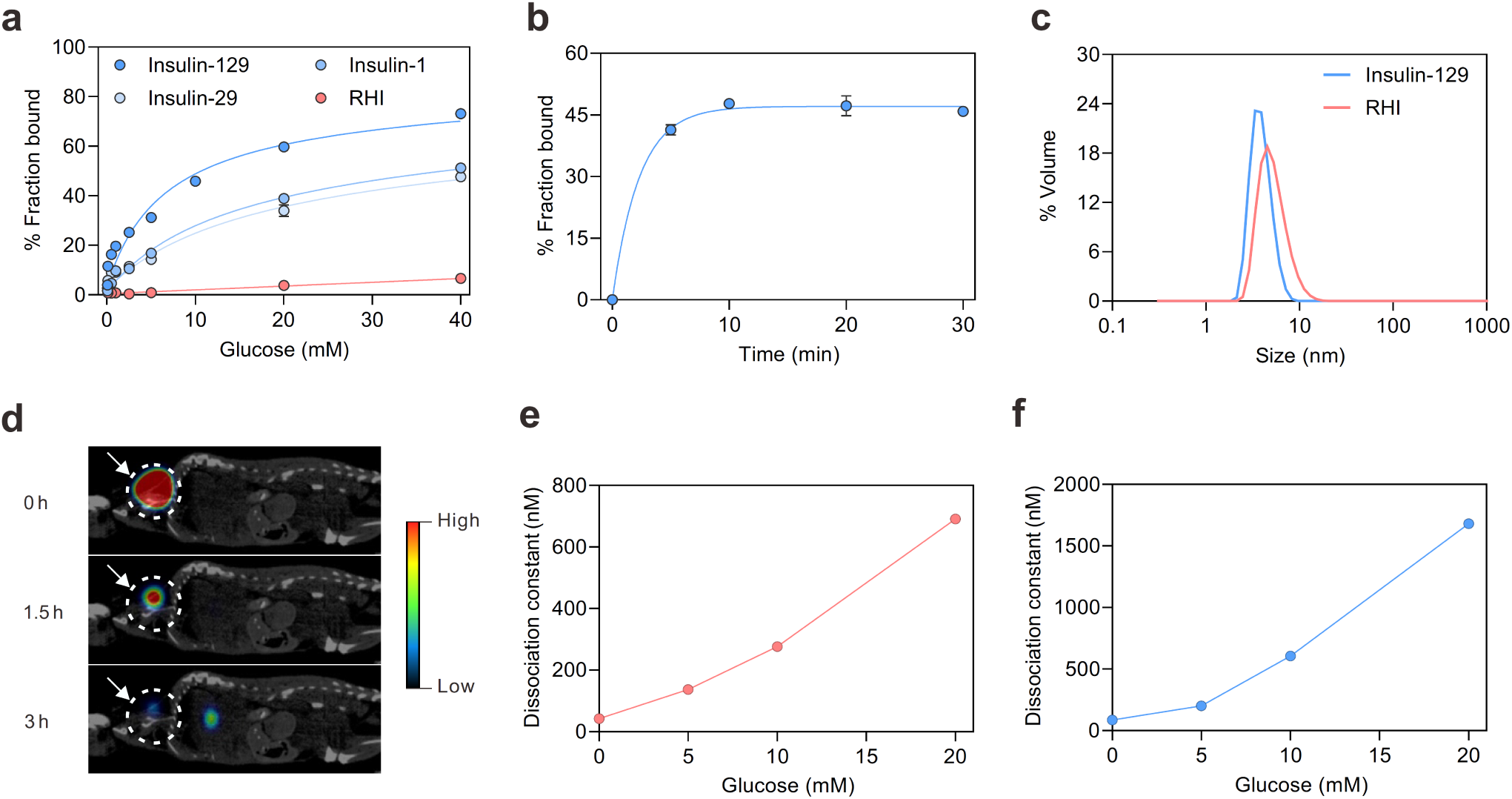
| Mechanism studies of insulin-129. **a,** *In vitro* glucose binding of insulin-129, insulin-1, insulin-29 and recombinant human insulin (RHI) analysed by native mass. Data points are means ± s.d. (*n* = 3). **b,** Glucose binding dynamics of insulin-129 under the glucose concentration of 10 mM. Data points are means ± s.d. (*n* = 3). **c,** Hydrodynamic sizes of RHI and insulin-129. The dynamic light scattering measurement by volume was performed at 25 °C in a 1× PBS buffer with a concentration of 3 mg/mL. **d**, Representative single photon emission computed tomography/computed tomography (SPECT/CT) images of type 1 diabetic mice after subcutaneous injection of insulin-129 (10 mg/kg). Insulin-129 was labelled by ^131^I using the chloramine T labelling method. The colour scale indicates the radiant efficiency from 0 to 0.25 percent of injected dose per (%ID/mL). **e,** The interactions of insulin-129 with IgG1-Fc under different glucose concentrations of 0, 5, 10 and 20 mM. **f,** The interactions of insulin-129 with nostrin under different glucose concentrations of 0, 5, 10 and 20 mM.

### Mechanism behind the long circulation and discovery of glucose-responsive binding to endogenous proteins

The prolonged duration can result from slow degradation, self-assembly to form large aggregates with reduced skin absorption rate, or binding to endogenous proteins in the circulatory system. When subjected to cathepsin D, a key contributor to the degradation of endogenous insulin, at a concentration of 62.5 mU/mL, insulin-129 exhibited a 3.2 times longer half-life than that of RHI (**Supplementary Fig. 2d**). However, this is not sufficient to explain the long circulation time of insulin-129. Then, dynamic laser scattering was used to study the self-assembly propensity of insulin-129 due to self-assembly’s important role in prolonging the treatment duration of long-acting insulin, like insulin detemir and insulin degludec ^34,35^. The hydrodynamic size of insulin-129 closely matched that of RHI across various concentrations, indicating no obvious formation of aggregates (**Fig. 2c and Supplementary Fig. 2e**). This result was consistent with the rapid skin absorption of ^131^I-labelled insulin-129 (**Fig. 2d**).

Then, the protein binding capability of insulin-129 was screened using a human protein microarray encompassing approximately 20,000 types of human being proteins. Surprisingly, IgG1-Fc and nostrin emerged as the principal binding partners. The binding of insulin-129 to IgG1-Fc and nostrin was studied using surface plasmon resonance (SPR). Mouse proteins were used in this test because the preliminary *in vivo* studies were executed in mice. In the absence of glucose, insulin-129 was found to bind to IgG1-Fc and nostrin with *K*_D_ values of 42.4 nM and 86.0 nM, respectively (**Fig. 2e,f and Supplementary Fig. 2f,g**). After adjusting glucose concentration to 5, 10, and 20 mM, *K*_D_ values of insulin-129 for IgG1-Fc were elevated to 137 nM, 276 nM, and 691 nM (**Fig. 2e and Supplementary Fig. 2f**), respectively. That is to say, a 16.3-fold and 5.0-fold increase were achieved, respectively, when the glucose concentration rose from 0 to 20 mM or from 5 to 20 mM glucose, the clinically relevant glucose range for people with diabetes. Similarly, the *K*_D_ of insulin-129 for nostrin increased to 201 nM, 605 nM, and 1680 nM, respectively (**Fig. 2f and Supplementary Fig. 2g**), indicating a 19.5-fold and 8.4-fold increase in *K*_D_ of insulin-129 for nostrin from 0 to 20 mM glucose or from 5 to 20 mM glucose.

In parallel, we performed computational simulations to explore the binding conformation of insulin-129 with mouse IgG1-Fc. Docking analyses revealed that the boron atoms introduced *via* FPBA modifications at residues A1 and B29 of insulin-129 form reversible covalent bonds with Glu170 and Trp260 of IgG-Fc. Moreover, Arg178 and Glu266 further stabilised this interaction through salt bridges and hydrogen bonding (**Supplementary Fig. 3a**). Similarly, the FPBA moieties of insulin-129 formed reversible covalent interactions with Ser33, Glu77, Glu118, Trp130, Ser186, and Glu218 residues of mouse nostrin. Beyond these covalent interactions, additional binding modes, including salt bridges, hydrogen bonds, and cation-π interactions, collectively reinforce the stability and specificity of the insulin-129–nostrin complex (**Supplementary Fig. 3b**). These docking-derived interaction patterns accord with previously reported protein interactions of boron-containing drugs, which typically involve reversible covalent adducts complemented by stabilising hydrogen-bond contacts ^36–40^. The engagement of FPBA in these interactions can provide insight into the glucose-responsive binding mechanism among insulin-129, IgG1-Fc, and nostrin.

### Systemic treatment efficacy study in type 1 diabetic mice

The dose-dependent therapeutic effects of insulin-129 were investigated in diabetic mice. After *s.c.* administration of insulin-129 at doses of 4, 8, and 12 mg/kg, the diabetic mice were kept normoglycaemic for 46, 142, and 190 hours, respectively (**Supplementary Fig. 2c and Fig. 3a**). Of note, the lowest average BG level of 76.0 ± 6.4 mg/dL was observed one hour after administration of insulin-129 at 12 mg/kg. No hypoglycaemia (BG < 70 mg/dL) in any mouse was observed. After further increasing the dose to 16, 20, and 24 mg/kg, the time in normoglycaemia gradually increased and reached a plateau (**Supplementary Fig. 4a**). By comparison, a single injection of insulin glargine (Lantus, U-100) at doses of 40 and 400 U/kg (equivalent to 1.46 and 14.6 mg/kg) only maintained normoglycaemia for 4 and 12 hours, respectively (**Fig. 3b**). Compared with insulin glargine, a single injection of insulin-129 extended normoglycaemia by approximately 47.5-folds. Notably, consecutive daily injections of insulin glargine (40 U/kg) exhibited poor glycaemic control (**Supplementary Fig. 4b,c**). The recently approved once-weekly insulin formulation, insulin icodec (Awiqli, U-700), was also evaluated in diabetic mice as a positive control. At a dose of 400 U/kg (equivalent to 15.3 mg/kg), insulin icodec maintained normoglycaemia for approximately 72 hours (**Supplementary Fig. 4d**), roughly one-third of the duration achieved by insulin-129 at a dose of 12 mg/kg.

**Fig. 3.**
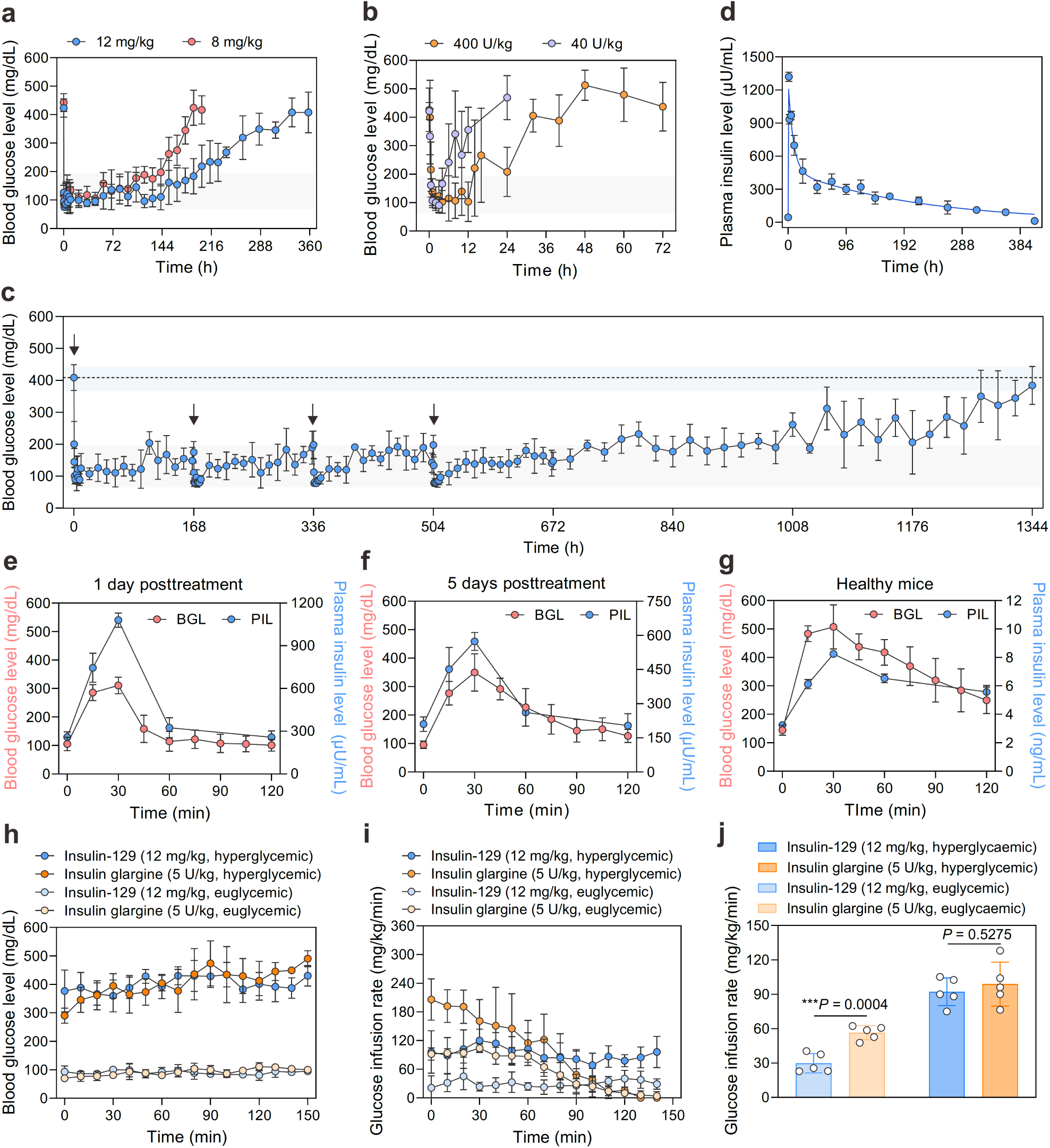
| Insulin-129 efficacy in type 1 diabetic mice. **a,** BG of diabetic mice treated with subcutaneously injected insulin-129 (8 and 12 mg/kg). Data points are means ± s.d. (*n* = 5). **b,** BG of diabetic mice treated with subcutaneously injected insulin glargine (Lantus, U-100, 40 and 400 U/kg). Data points are means ± s.d. (*n* = 5). **c,** BG of diabetic mice treated four times with subcutaneously injected insulin-129 at 0, 168, 336 and 504 h. The first loading dose of insulin-129 was 10 mg/kg. The following three doses were 4.8 mg/kg. Data points are means ± s.d. (*n* = 5). **d,** Pharmacokinetics of insulin-129 after a single subcutaneous injection in type 1 diabetic mice (12 mg/kg). Data points are means ± s.d. (*n* = 5). **e,f,** Hyperglycaemia-triggered insulin release after intraperitoneal glucose injection (3 g/kg) at one day (**e**) and five days (**f**) after treatment of insulin-129 at a dose of 12 mg/kg. Data points are means ± s.d. (*n* = 5). **g,** Hyperglycaemia-triggered insulin release by intraperitoneal glucose injection (3 g/kg) in healthy mice. Data points are means ± s.d. (*n* = 5). **h,i,** Isoglycaemic clamp experiments, during which the steady-state BG concentration (**h**) and the corresponding glucose infusion rate (**i**) were monitored in healthy mice (euglycaemic) or type 1 diabetic mice (hyperglycaemic). The glucose infusion was carried out one day after the administration of insulin-129 (12 mg/kg) or immediately after the injection of insulin glargine (5 U/kg). Data points are means ± s.d. (*n* = 5). **j,** Average glucose infusion rate of (**i**). Data points are means ± s.d. (*n* = 5) and individual points. Unpaired two-tailed Student’s *t*-test was used for statistical analysis. \*\*\**P* < 0.001. All *P* value analyses were performed on GraphPad Prism (version 10.0).

The long-term treatment efficacy of insulin-129 in BG regulation was then assessed through four sequential weekly injections. The initial dose was set to be 10 mg/kg (a loading dose), followed by three subsequent weekly-administered normal doses of 4.8 mg/kg. Each injection could maintain normoglycaemia for one week, with only a rapid BG drop to 89.6 ± 13.1 mg/dL, 79.6 ± 12.4 mg/dL, 77.8 ± 5.4 mg/dL, 78.8 ± 8.8 mg/dL in the first few hours after the first to fourth injections. Most notably, BG stabilised at approximately 150 mg/dL one day after the fourth injection (**Fig. 3c**). After the fourth injection, insulin-129 remained effective for more than 800 hours before BG returned to the original BG level before the first injection.

Then, the pharmacokinetics of insulin-129 was studied. Diabetic mice were *s.c.* administered insulin-129 at 12 mg/kg, with blood sampling (20 μL) at different intervals. The insulin-129 levels were measured using enzyme-linked immunosorbent assay (ELISA) kits. Before treatment, insulin-129 was virtually undetectable in the plasma of diabetic mice (**Supplementary Fig. 4e**). Following *s.c.* injection, the average plasma insulin-129 level (PIL) peaked at 1320.0 ± 41.3 μU/mL within one hour and then decreased very slowly. The pharmacokinetic curve was well-fitted to a two-compartment model. Based on the curve, the t_1/2*α*_ and t_1/2*β*_ of insulin-129 were calculated to be 5.4 ± 1.3 hours and 147.8 ± 16.8 hours (**Fig. 3d**). Further analysis following intravenous dosing in healthy mice revealed an average t_1/2*β*_ of 112.5 ± 17.4 hours (**Supplementary Fig. 4f**).

### Glucose tolerance test and glucose-responsive insulin release study in diabetic mice

Glucose tolerance tests were conducted to evaluate the BG-regulating efficacy of insulin-129. Mice received insulin-129 at 12 mg/kg and an intraperitoneal glucose injection at 1.5 g/kg one and five days post-treatment. After glucose administration, BG rose from 96.8 ± 19.7 mg/dL (day 1) and 163.1 ± 27.7 mg/dL (day 5) to a peaking BG of 194.8 ± 31.9 mg/dL and 310.3 ± 44.2 mg/dL within 15 minutes, and then gradually returned to original levels (**Supplementary Fig. 4g,h**). In contrast, healthy mice displayed steeper BG peaks and failed to return to baseline within 120 minutes. The area under the curve (AUC) for the BG-time curve in insulin-129-treated mice was significantly smaller than that of healthy mice for both tests performed on day one and day five (**Supplementary Fig. 4i**).

BG-stimulated insulin release was further explored in insulin-129-treated mice. Glucose dose was increased to 3 g/kg to induce marked BG spikes at one and five days posttreatment. After each injection, BG spikes were all accompanied by insulin spikes synchronously. One day after insulin-129 treatment, a distinct BG spike of 310.7 ± 29.2 mg/dL was observed at 30 minutes following glucose administrations (**Fig. 3e**). Subsequently, BG levels rapidly dropped to be below 200 mg/dL by 45 minutes and continued to decline gradually over time (**Fig. 3e**). Corresponding to the rise in BG, PIL increased approximately 4.2-fold, from 258.6 ± 37.6 μU/mL to a peak of 1080.4 ± 49.6 μU/mL, before gradually declining to 257.3 ± 45.2 μU/mL in tandem with the BG reduction. As for five days after insulin-129 administration, PIL peaked at 573.7 ± 40.0 μU/mL, approximately 2.2 times the initial level of 210.2 ± 31.7 μU/mL, and declined to 204.3 ± 52.0 μU/mL within 120 minutes (**Fig. 3f**). In contrast, insulin glargine (Lantus, U-100) did not display BG-correlated changes (**Supplementary Fig. 4j,k**). As another control, healthy mice showed a 2.5-fold plasma insulin level increase in response to the same glucose challenge but were unable to maintain normoglycaemia at 2 hours post-injection (**Fig. 3g**).

### Isoglycaemic clamp studies in mice

To further assess the *in vivo* glucose-responsive properties of insulin-129, particularly its hypoglycaemia-mitigating capacity, we performed isoglycaemic clamp experiments in both healthy and diabetic mice. For the isoglycaemic clamp at hyperglycaemia in diabetic mice, glucose was infused *via* venous catheters to maintain blood glucose at 400 mg/dL, which was near its original level (**Fig. 3h**). This infusion was initiated either one day after administering insulin-129 (12 mg/kg) or immediately following insulin glargine injection (5 U/kg). We also conducted a euglycaemic clamp in healthy mice by infusing glucose to maintain BG at approximately 100 mg/dL (**Fig. 3h**). The AUC for the BG-time curve was similar between insulin-129 and insulin glargine groups at both hyperglycaemia and normoglycaemia, indicating a comparable average BG level (**Supplementary Fig. 4l**). During isoglycaemic clamp study at hyperglycaemia, insulin-129 and insulin glargine groups exhibited comparable mean glucose infusion rates (**Fig. 3i,j**). However, during the isoglycaemic clamp study at normoglycaemia in healthy mice, the average glucose infusion rate in the insulin-129 group required only one-half of that in the insulin glargine group (**Fig. 3i,j**). This significantly reduced glucose clearance of insulin-129 at reduced BG conferred a protective effect against hypoglycaemia.

### Treatment efficacy and glucose tolerance study of insulin-129 in minipigs

The treatment efficacy of insulin-129 was further evaluated in three type 1 diabetic minipigs induced chemically (designated Pig-Ⅰ, Pig-Ⅱ, and Pig-Ⅲ). BG levels of diabetic minipigs were monitored by the CGMS (SIBIONICS) with an effective range of 39.6 to 450.0 mg/dL. The sensor of CGMS was stuck onto the inner thigh of minipigs. The minipigs were fed twice every day. The average BG levels of these three diabetic minipigs before treatment were all above 250 mg/dL. As the positive control, insulin glargine (Lantus, U-100) was given at 0.25, 0.33, and 0.48 U/kg to Pig-Ⅰ, Pig-Ⅱ and Pig-Ⅲ, respectively. A single dose of insulin glargine regulated BG in the normal range for less than 12 hours in Pig-Ⅰ and Pig-ⅡI and less than 24 hours in Pig-II (**Fig. 4a**). After a two-day gap without treatment, insulin-129 was *s.c.* administrated at doses of 0.16, 0.36 and 0.63 mg/kg to Pig-Ⅰ, Pig-Ⅱ, and Pig-Ⅲ, respectively. Following injections, all minipigs exhibited consistent BG control over the course of one week, with the 9-day average daily BG levels stabilizing within the normal range (**Fig. 4a and Supplementary Fig. 5a**), comparable to BG control observed in the healthy pig (**Supplementary Fig. 5b**).. The BG level in the insulin-129-treated group remained within the target range (70 to 200 mg/dL) for 73.0% of the time, whereas the Lantus-treated group achieved only 42.9% (**Supplementary Fig. 5c**).

**Fig. 4.**
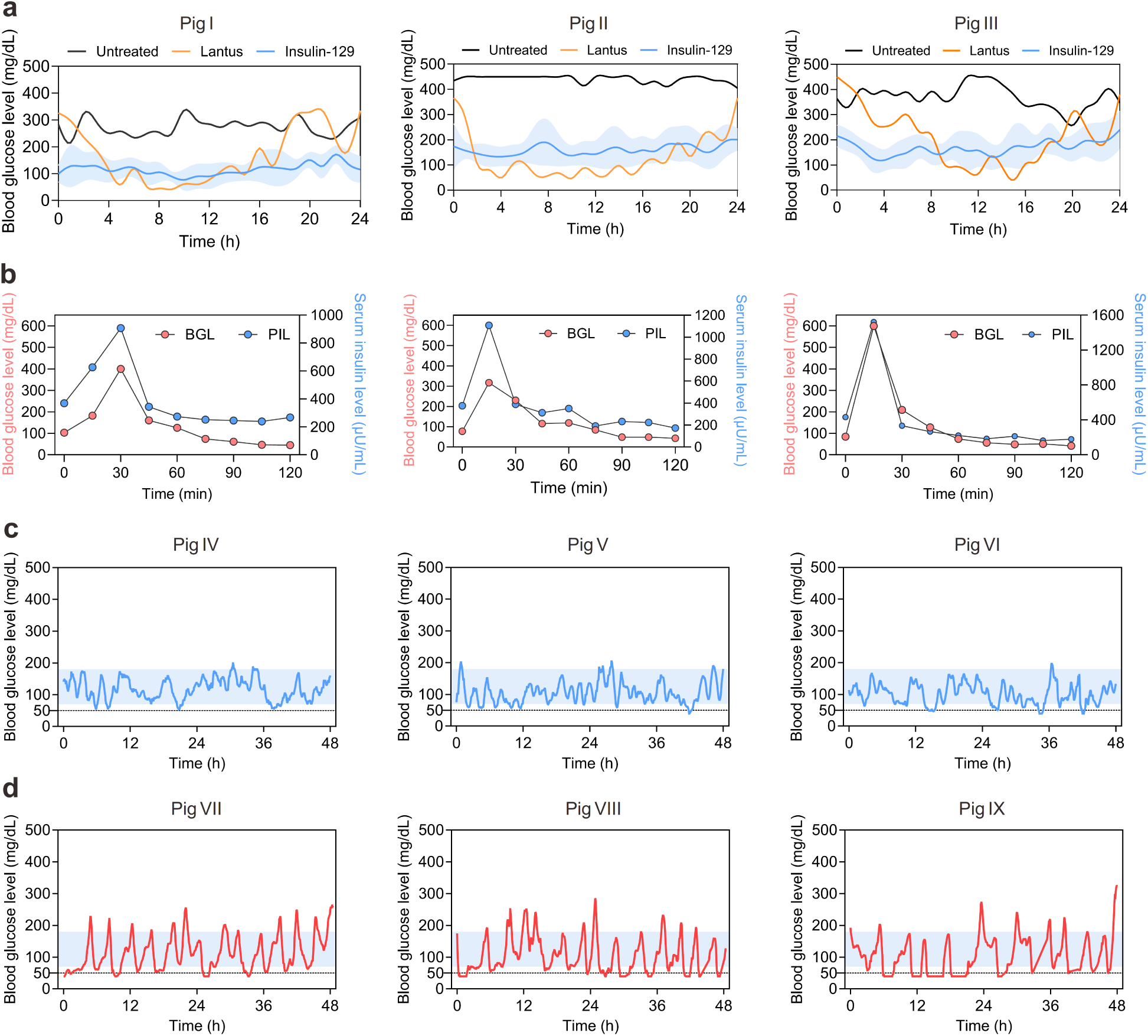
| Evaluation of insulin-129 in type 1 diabetic minipigs and its use in a dual-closed-loop fully automated insulin delivery system. **a,** BG levels in type 1 diabetic minipigs without treatment (black line), with subcutaneous insulin glargine at doses of 0.25, 0.33, and 0.48 U/kg for Pig I, Pig II, and Pig III (orange line), and as 9-day averages following a single injection of insulin-129 at 0.16, 0.36, and 0.63 mg/kg for Pig I, Pig II, and Pig III (blue line, shadows represent standard deviation ranges). All curves were smoothed using the cubic spline method by GraphPad Prism (version 10.0). **b,** Hyperglycaemia-triggered insulin release. Dextrose (10 wt%) solution was infused at a dose of 0.75 g/kg at a rate of 1 L/h until the BG of the minipigs was above 300 mg/dL while the minipigs were kept anaesthetised using isoflurane. BG was measured using a glucose meter. Each figure is associated with data collected from one minipig. **c,** Representative BG of type 1 diabetic minipigs treated with insulin-129 using a dual-closed-loop fully automated insulin delivery system. If two consecutive BG were more than 150 mg/dL, insulin-129 was injected at 0.2, 0.3, and 0.2 mg per time, respectively, with an interval of more than 2 h between each dose. **d,** BG of type 1 diabetic minipigs treated with insulin lispro using a fully automated insulin delivery system. If two consecutive BG were more than 150 mg/dL, insulin lispro was injected at 0.24, 0.14, and 0.12 mg per time, respectively, with an interval of more than 2 h between each dose.

The pharmacokinetics of insulin-129 were investigated in all diabetic minipigs by collecting blood samples after administering insulin-129 at doses of 0.16 mg/kg to Pig-I, 0.22 mg/kg to Pig-II, and 0.32 mg/kg to Pig-III, respectively. Insulin-129 was undetectable prior to injection (**Supplementary Fig. 5d**). Following injections, all three minipigs exhibited an obvious peak in serum insulin-129 level at 6 hours, which gradually decreased over time to baseline after 288 hours (**Supplementary Fig. 5e**). The half-life of insulin-129 in diabetic minipigs was determined to be 127.5 ± 36.6 hours using the two-compartmental pharmacokinetic model.

Glucose tolerance tests were studied one day after administering insulin-129 by infusing glucose intravenously until BG exceeded 300 mg/dL. The minipigs were kept anaesthetised by isoflurane breath. Within 30 minutes, BG in Pigs Ⅰ, Ⅱ, and Ⅲ rose sharply to 399.6, 316.8, and 599.4 mg/dL, respectively, then fell below 200 mg/dL by 45 minutes (**Fig. 4b**). In parallel with these BG fluctuations, the serum insulin-129 level rose by approximately threefold from baseline values (369.1, 375.5, and 430.6 μU/mL for Pigs I, II, and III, respectively) to their corresponding peaks (907.3, 1108.0, and 1520.5 μU/mL). This surge effectively facilitated the rapid normalisation of BG levels, achieving euglycaemia by the next measurement following the cessation of glucose infusion. At the end of this experiment, the BG decreased to a level lower than the initial level probably due to anaesthetisation.

### Application of insulin-129 in a dual-closed-loop fully automated insulin delivery system

To date, no fully automated insulin delivery system without operator input has been approved. To address this, we replaced standard fast-acting insulin, insulin lispro with insulin-129, because of the features of rapid absorption and sustained, glucose-responsive release. The resulting platform integrated a CGMS, a host device (Android APS for open-source algorithm customization), a custom algorithm, and an insulin pump, forming a dual-closed-loop fully automated insulin delivery system (DF-AID). The same algorithm was set up to adapt to insulin-129 and insulin lispro for AID. In DF-AID, the initial parameters of the delivery algorithm contained a minimal administration interval (set at 2 h), dose (set to 0.2, 0.3, or 0.2 mg, dependent on the minipigs), and blood glucose at infusion (set at 150 mg/dL). For SC-AID, the delivery algorithm was similar to that of DF-AID, but with slightly altered insulin lispro doses (0.24, 0.14, or 0.12 mg, for another three minipigs). During the operation of both systems, no human intervention or parameter input was allowed. BG points were recorded by the CGMS and updated every five minutes. BG and delivery doses were monitored and collected continuously for 48 h (**Fig. 4c,d**), and the implementation rates of both systems were > 95%.

The BG of diabetic minipigs treated with insulin-129 were stable, with BG levels rarely dipping below 50 mg/dL (specific values of 0, 1.0, and 4.2%, respectively and an average of 1.7 ± 2.2%), which is considered severe hypoglycaemia in humans. The AUC of the BG-time curve had no significant difference between the treatment of DF-AID and SC-AID (**Supplementary Fig. 5f**). However, the coefficient of glucose variation for minipigs treated with DF-AID (30.5 ± 2.1%) was lower than that of SC-AID (54.7 ± 3.5%), indicating a much reduced BG fluctuation for DF-AID (**Supplementary Fig. 5g**). The safety and efficacy of BG management were further evaluated in terms of the percentage of glucose time in range (TIR, 70-180 mg/dL), time above range (TAR, >180 mg/dL), and time below range (TBR, < 70 mg/dL). The TIR of DF-AID was 85.6 ± 4.1%, significantly higher than that of SC-AID (56.0 ± 2.6%). Of note, the TIR of DF-AID using insulin-129 was similar to that of hybrid systems ^41,42^. The TBR of DF-AID (12.4 ± 4.5%) was lower than that of SC-AID (33.7 ± 3.4%), demonstrating a reduced occurrence of hypoglycaemia. Notably, the TAR observed with DF-AID (2.0 ± 1.1%) was markedly lower than that of SC-AID (10.3 ± 2.1%) (**Supplementary Fig. 5h**).

### Disposition study in mice

Cy5-labelled insulin-129 was used to visualise the disposition of insulin-129 in diabetic mice, with free Cy5 or Cy5-labelled RHI used as controls. After *s.c.* injection in mice, the fluorescence of free Cy5 disappeared most rapidly through the kidney, while the fluorescence of Cy5-labelled RHI persisted for 24 hours (**Supplementary Fig. 6a**). Within one hour of injection of Cy5-labelled RHI, prominent fluorescence signals in the kidney, lung, and liver were observed. The high fluorescence intensity in the bladder indicated rapid excretion through the urinary pathway (**Supplementary Fig. 6b**). The signal in these organs attenuated over time until declining below the background threshold. Cy5-labelled insulin-129 had a similar fluorescence biodistribution to Cy5-labelled RHI (**Fig. 5a,b**); however, the signal intensity of Cy5-labelled insulin-129 in the lung showed a sustained increase until reaching its maximum at 24 hours, followed by a gradual decrease till 168 hours post-injection. The detailed tissue distribution of insulin-129 was further analysed. Cy5-labelled insulin-129 exhibited a high level of vascular association within the major organs harvested three days after subcutaneous injection (**Fig. 5c,d**), consistent with the vascular distribution of FcRn and nostrin ^43,44^.

**Fig. 5.**
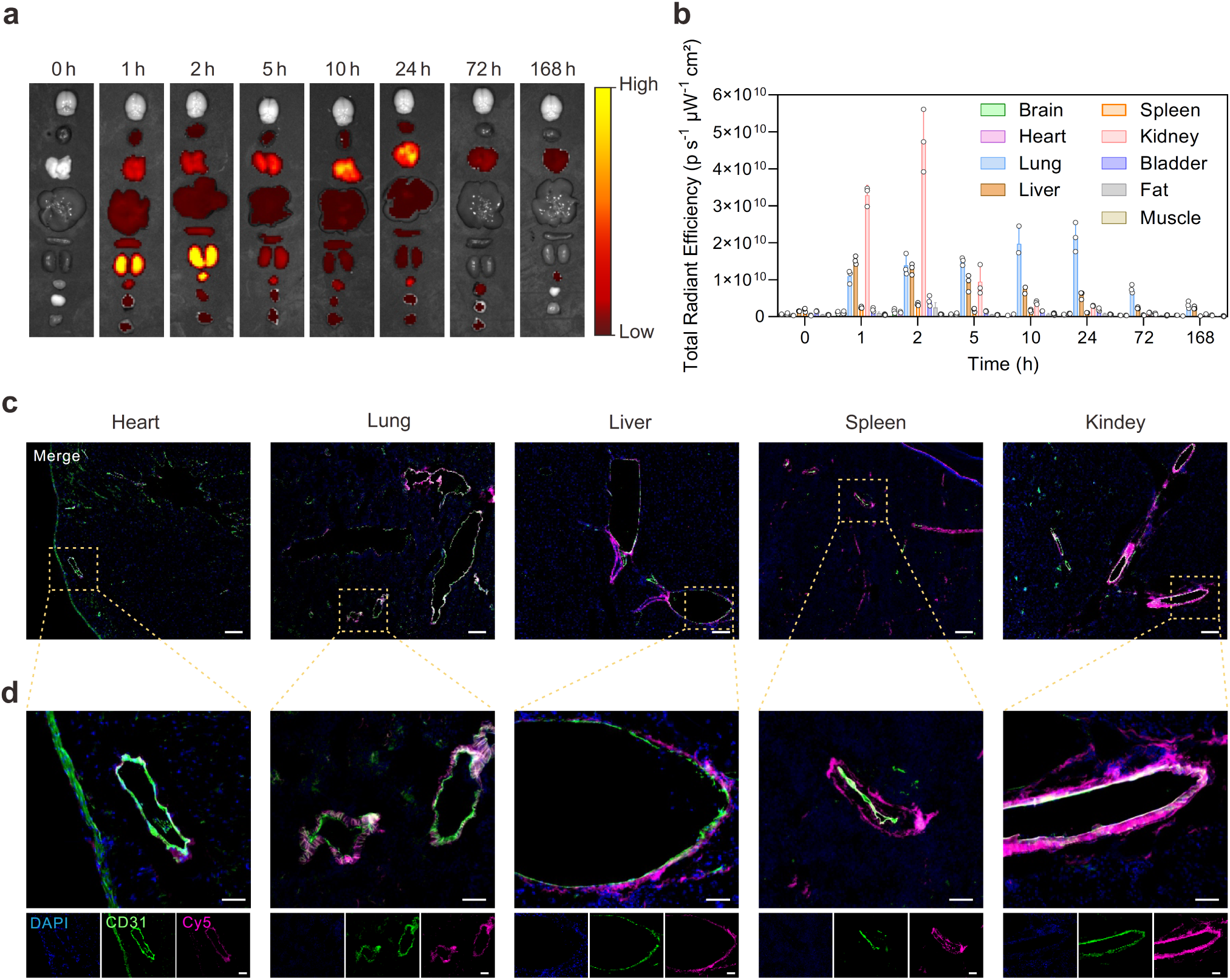
| Disposition and biosafety of insulin-129. **a,** Representative *ex vivo* fluorescence imaging of type 1 diabetic mice after subcutaneous injection of Cy5-labelled insulin-129 (10 mg/kg). The major organs or tissues (from top to bottom) included the brain, heart, lung, liver, spleen, kidney, bladder, fat and skeleton muscle. The colour scale indicates the radiant efficiency from 1.00 × 10^8^ to 3.00 × 10^9^ photons per s per cm^2^ per sr. **b,** Fluorescence intensity-time profiles of major organs or tissues in (**a**). Data points are means ± s.d. (*n* = 3). **c,** Representative fluorescence images of the heart, lung, liver, spleen and kidney sections at 3 days after the subcutaneous administration of Cy5-labelled insulin-129 at a dose of 10 mg/kg (purple). Cell nuclei were stained with DAPI (blue) and the blood vessels were stained with anti-mouse CD31 monoclonal antibody (green). Scale bars, 200 μm. **d,** The detailed manually selected images of the blood vessels of the main organs in (**c**). Scale bars, 50 μm.

### Biosafety evaluation of insulin-129 in mice

We evaluated the biosafety of insulin-129 in mice by analysing key biochemical parameters, blood cell counts, and tissue morphology. Mice received either a single *s.c.* injection of insulin-129 (12 mg/kg) or four weekly injections (10 mg/kg for the first dose and 4.8 mg/kg for the subsequent three doses). Blood samples were collected on days 7 and 28, respectively, to assess its safety profile.

No significant differences between the groups were observed by blood routine tests. Of note, white blood cell and lymphocyte levels in insulin-129-treated mice were more closely aligned with those of healthy controls compared to untreated diabetic mice (**Supplementary Fig. 6c**). Notably, long-term treatment with insulin-129 progressively brought blood routine test parameters closer to those observed in healthy mice. In untreated diabetic mice, all tested biochemical markers were significantly elevated compared to healthy controls. However, following treatment with insulin-129, these markers returned to levels indistinguishable from those of healthy mice, except albumin values, which remained elevated after one month of continuous treatments (**Supplementary Fig. 6d**). In addition, histomorphological analysis of major organs by H&E staining one week post-injection revealed no apparent pathological alterations, further supporting the favourable safety profile of insulin-129 (**Supplementary Fig. 6e,f**).

## Conclusions

This study presented an ultra-long acting GRI named insulin-129 that can dynamically adjust its blood concentration in synchrony with BG changes, a feature critical for real-time BG regulation but has been rarely achieved. Insulin-129 bound to blood circulating proteins and endothelial proteins like IgG1-Fc and nostrin tightly. Of note, insulin-129 also exhibited similar strong binding capability to human IgG1-Fc and nostrin, with *K*_D_ values of 36.7 nM and 241 nM (**Supplementary Fig. 7**). This finding indicates that this glucose-responsive binding mechanism is preserved in humans and highlights insulin-129’s clinical translational potential. After s.c. injection, insulin-129 formed a dynamic reservoir in the blood and along the systemic vascular system, exhibited ultralong circulatory time, and sustained normoglycaemia for more than one week in type 1 diabetic mice and minipigs. Insulin-129 also bound dynamically to glucose with rapid kinetics and exhibited a physiologically relevant *K*_D_ value. As a result, glucose can modulate insulin-129’s binding strength toward the endogenous proteins like IgG1-Fc and nostrin. Once BG increased, a synchronised elevation of circulating insulin-129 levels can quickly normalise the elevated glucose to the normal range, a response likely derived from the detachment of insulin-129 from endogenous proteins. This closed-loop insulin delivery behaviour also contributed to the minimal hypoglycaemia risk of insulin-129 during treatment. Eventually, a fully automated insulin delivery system was established on insulin-129, which exhibited negligible hypoglycaemia risk and a glucose time in range higher than the clinical target.

Overall, this study adopts a unique mode of real-time glucose detection and response through glucose-dependent dynamic protein-protein interactions. It provides a novel paradigm of prolonged duration of insulin and realises glucose sensitivity by harnessing endogenous proteins other than albumin. Importantly, the ability to rapidly adapt to glycaemic fluctuations highlights the translational relevance of this approach, as real-time responsiveness is critical for minimising hypo- and hyperglycaemic excursions and for meeting clinical requirements in GRIs ^45^.

## Acknowledgements

This work was supported by grants from National Key R&D Program of China (2022YFE0202200), the Key R&D Program of Zhejiang Province (2024C03085, J.W.), the Innovation and Technology Fund (Mainland-Hong Kong Joint Funding Scheme, MHP/053/21), the Shenzhen-Hong Kong-Macau Technology Research Program (SGDX20210823103537034), National Natural Science Foundation of China (32471374, J.W.), the Zhejiang University’s Start-Up Packages, the Starry Night Science Fund at Shanghai Institute for Advanced Study of Zhejiang University (SN-ZJU-SIAS-009, J.W.), the Postdoctoral Fund of Zhejiang Province (ZJ2023044, J.X.) and the Postdoctoral Fellowship Program of CPSF (GZB20240669, L.L.). We thank Guizhen Zhu in the Center of Cryo-Electron Microscopy (CCEM), Zhejiang University for their technical assistance in Scanning Electron Microscopy. We acknowledge Long Shi and Shiwen Zhou at the Chemistry Instrument Center and the Consortium for Advanced Technologies of Mass Spectrometry at Zhejiang University for the mass spectrometry instrument support. We acknowledge Deyue Xu, Maolong Zhang, and Shenglong Xiong (Animal Center, Zhejiang University) for taking care of the minipigs.

## Author contributions

Y.Z. and J.Z. contributed equally to this work. Z.G., J.W., and Y.Z. conceived and designed the study. Y.Z., J.Z., K.J., S.M., H.Z., J.L., X.H., X.Z., K.X., J.X., L.L., J.Y., C.J., and X.L. conducted experiments and obtained related data. C.S. and J.Y. gave experimental operation teaching and theoretical guidance on isotope labelling. M.Z. gave experimental operation teaching and theoretical guidance on native mass analysis. Y.W., Y.Z., and Z.G. designed and drew the schematic of this article. Y.Z., J.W., J.Z., J.B.B., P.L., S.Z., and Z.G. analysed the data and wrote the paper.

## Materials and Methods

### Materials

All materials were reagent grade or better. 4-carboxy-3-fluorophenylboronic acid was purchased from Aladdin. Recombinant human insulin was purchased from Solarbio (catalogue no. I8830). Recombinant mouse and human nostrin were purchased from Lead Biotech. Recombinant mouse and human immunoglobulin heavy constant gamma 1 were purchased from Medlife. Cathepsin D was purchased from Sigma-Aldrich. (4-(((2,5-dioxopyrrolidin-1-yl)oxy)carbonyl)-3-fluorophenyl)boronic acid (FPBA-NHS) was synthesised as reported ^46^.

### Preparation of insulin-129

FPBA-NHS (362.5 mg, 1.29 mmol) was dissolved in DMSO (10 mL) in a 25 mL round-bottom flask under magnetic stirring. At the same time, recombinant human insulin (RHI) (5 g, 0.86 mmol) and *N*, *N*-diisopropylethylamine (180.0 μL, 1.03 mmol) were mixed in anhydrous DMSO (50 mL). FPBA-NHS solution was added dropwise into the RHI solution, followed by one hour of stirring at room temperature. Then, the mixture was poured into a mixture of acetone and methyl tert-butyl ether (4:1, 500 mL) to afford a white precipitate. The suspension was stirred for 30 min, filtered, and washed with tetrahydrofuran (3 times, 200 mL each time). The solid was further purified by preparative HPLC to yield insulin-129 (750.0 mg, 98.4% purity, 14.3% yield).

Other insulin analogues were prepared in similar ways.

### Preparation of Cy5-labelled insulin-129

Sulfo-Cy5-NHS (25 μL, 25 mg/mL in DMSO, 0.82 μmol) was added to a solution of insulin-129 (50 mg, 8.22 μmol) in DMSO (2 mL) containing *N*, *N*-diisopropylethylamine (6.25 μL, 35.8 μmol). The mixture was stirred at room temperature for 15 min. The solution was added to a vigorously stirred mixture of acetone: methyl tert-butyl ether (4:1, 20 mL), affording a blue precipitate. The suspension was stirred for 10 min, centrifuged (9500 × g, 5 min), and washed with acetone (3 times, 10 mL each time). The solid was lyophilised to yield Cy5-labelled insulin-129 (50.8 mg, 88.3% yield).

### Preparation of Cy5-labelled RHI

Sulfo-Cy5-NHS (100 μL, 25 mg/mL in DMSO, 3.28 μmol) was added to a solution of insulin (10 mg, 1.64 μmol) in DMSO (400 μL) containing *N*, *N*-diisopropylethylamine (1.25 μL, 7.16 μmol). The mixture was stirred at room temperature for 15 min. The solution was added to a vigorously stirred mixture of acetone: methyl tert-butyl ether (4:1, 5 mL), affording a blue precipitate. The suspension was stirred for 10 min, centrifuged (9500 × g, 5 min), and washed with acetone (3 times, 2 mL each time). The solid was lyophilised to yield Cy5-labelled RHI (11.2 mg, 89.6% yield).

### Preparation of ^131^I-labelled insulin-129

Insulin-129 in PBS (1 mg/mL, 20 µL) was mixed with chloramine-T (1 mg/mL, 30 µL) and Na^131^I (5 mCi). The mixture was incubated at 30°C for 15 min to afford the product.

### Characterization of insulin-129

The purity of insulin-129 and other insulin analogues was analysed using high-performance liquid chromatography (HPLC, Waters E2695, USA). The eluent was acetonitrile with 0.1% trifluoroacetic acid and water (50:50, v/v). The flow rate was set at 1.0 mL/min. The absorbance was measured at 280 nm. The insulin secondary structure was characterized through a circular dichroism (CD) spectrometer (J-1500-150ST, JASCO, Japan). All insulin analogues were tested at a concentration of 0.2 mg/mL in PBS (pH 7.4).

### Hydrodynamic diameters of insulin-129 and RHI

Insulin-129 and RHI were dissolved in PBS (pH 7.4) at concentrations of 0.25, 0.5, 1 and 3 mg/mL. The hydrodynamic diameters were tested using a dynamic light scattering set-up (Zetasizer Nano ZS; Malvern Instruments)

### Mass spectrometry analysis

MS experiments under denaturing conditions were conducted on a Thermo Orbitrap Eclipse mass spectrometer with the Nanospray Flex source (San Jose, CA, USA). Static nanoelectrospray was achieved *via* in-house made borosilicate capillaries (catalogue BF100-78-10, Sutter Instrument, Novato, CA, USA). Protein samples were diluted at 0.1 mg/mL in 50% acetonitrile, 0.1% formic acid. Both intact and reduced proteins (treated with tris(2-carboxyethyl)phosphine, TCEP) were analysed. The application mode was set at intact protein mode, with the electrospray ionisation needle held at 1.0 kV. The heated capillary was set at 250°C. The MS system was calibrated using a Thermo positive calibration mix. All MS/MS experiments were conducted with a data-dependent acquisition method in HCD cells at 35 eV collision energy with N_2_ as collision gas. Source fragmentation was used in all MS and MS/MS experiments. Raw data were processed with TopPIC ^47^, and results were examined manually to locate the modification sites.

### Activity evaluation of insulin-129

The HepG2 cell line (acquired from the National Collection of Authenticated Cell Cultures, Shanghai, China) was propagated in Dulbecco’s modified Eagle’s medium (DMEM) supplemented with 10% fetal bovine serum (FBS) and 1% penicillin-streptomycin. Once the culture reached approximately 80–90% confluence, the cells were transferred to a serum-free medium for 12 h. They were then washed and incubated with a medium containing RHI or insulin-129 at concentrations of 0, 0.1, 1, 5, 10, 50, and 100 nM for 10 min (*n* = 3).To prepare lysates, cells were harvested in ice-cold RIPA buffer (Beyotime) containing protease and phosphatase inhibitors (Meilunbio), and the resulting lysate was centrifuged at 16000 × g for 10 min at 4°C. The protein content of the supernatant was determined by a BCA assay (Beyotime). Equal amounts of protein were combined with 5 × Laemmli loading buffer (Bio-Rad) containing 2.5% β-mercaptoethanol and heated to induce denaturation. Proteins were separated using 4-20% SDS–PAGE and transferred onto 0.45 µm polyvinylidene difluoride (PVDF) membranes (Sigma-Aldrich). After blocking with 5% bovine serum albumin (BSA) in phosphate-buffered saline containing 0.1% Tween-20, the membranes were incubated with primary antibodies (anti-phospho-Akt, Cell Signaling Technology, catalogue no. 4060, 1:2,000; anti-Akt, Cell Signaling Technology, catalogue no. 4691, 1:1,000; and anti-β-actin, Servicebio, catalogue no. GB11001, 1:1,000). Following primary incubation, membranes were treated with horseradish peroxidase-conjugated secondary antibodies (Hangzhou Fude, catalogue no. FDR007, 1:10,000) and visualised using an ECL Western Blotting Substrate (Thermo Fisher Scientific), following the manufacturer’s instructions.

### *In vitro* glucose-binding of insulin-129

Prior to glucose addition, insulin-129 (1.0 mg/mL) was first subjected to buffer exchange into 100 mM ammonium acetate (NH_4_Ac) at pH 7.4 using Amicon ultra centrifugal devices equipped with a 3,000 Da molecular weight cutoff filter. Glucose solutions, ranging from 0.05 to 40 mM (0.1, 0.2, 1, 2, 5, 10, 20, 40, and 80 mM), were prepared in the same NH_4_Ac solution. Each glucose solution was subsequently mixed with the buffer-exchanged insulin-129 at equal volumes. All mixtures were incubated for 30 min at 37℃. Subsequently, direct infusion was carried out using a nanoelectrospray source on a Waters Synapt XS mass spectrometry system (Manchester, United Kingdom). Analysis was performed in positive ionisation mode using the same static nanoelectrospray setup described earlier but under native conditions. The glucose-bound fraction was calculated by dividing the intensity of the glucose-insulin-129 complex by the TIC intensity of total insulin-129 using the formula of ((M + 1 glucose) + (M + 2 glucose))/(M + (M + 1 glucose) + (M + 2 glucose)). The peak intensities were obtained from the most abundant isotope peaks of the centroid spectra in MassLynx. The percentage of the glucose-bound fraction was plotted against glucose concentration (mM) using GraphPad Prism (version 10.0). Insulin-1, insulin-29, and RHI were tested similarly. The apparent equilibrium *K*_D_ for glucose was derived from the binding curves by fitting the data to either a one-site total and nonspecific binding model. The binding curve of RHI was fitted by a one-site total binding model.

The glucose binding dynamics toward insulin-129 were carried out using the same system. 10 mM glucose solution in the presence of insulin-129 (0.5 mg/mL)was prepared in 100 mM NH_4_Ac. For the zero point, insulin-129 solution (0.5 mg/mL in 100 mM NH₄Ac) was tested in the absence of glucose. As no glucose binding was detected, the bound fraction was assigned a value of zero. Timing was initiated immediately upon sample preparation, with measurements conducted at 5, 10, 20, and 30-minute intervals. It is important to note that these time points represent the duration from sample mixing to the application of voltage in the mass spectrometry analysis. The glucose binding dynamics curve was fitted by one-phase exponential association model using GraphPad Prism (version 10.0).

### Screening for binding proteins of insulin-129 using a protein chip

Protein binding microarray chips comprised of about 20000 individual human glutathione S-transferase (GST)- and His-tagged full-length proteins were obtained from the Johns Hopkins Medical Institutions Protein Microarray Core (CDI Laboratories, Inc). The microarray proteome analysis was performed according to the procedure detailed below, and the experiments and data processing were performed by Wayen Biotechnology Company (Shanghai, China).

Briefly, HuProt proteome microarrays were blocked with blocking buffer (5% BSA in 1×TBST, pH 7.5) for 1.5 hours at room temperature, followed by washing (1×TBST) for 5 minutes. Biotin-insulin-129 in blocking buffer (10 µmol/L) was then incubated with the blocked proteome microarrays at room temperature for 1 hour. Thereafter, the microarrays were washed 3 times with TBST, for 5 minutes each time, and Cy5-Streptavidin was added (1:1000 dilution). Following incubation with Cy5-Streptavidin, microarrays were washed with TBST (3 times, 5 minutes each time) again. Finally, they were spun for 2 minutes and then scanned with an Axon GenePix 4000B. The GenePix Pro 6.0 software (Axon Instruments) extracted data from the recorded microarray images. Protein spots with a signal intensity indicator Z-Score > 2.8 in biotin-insulin-129-treated microarrays and < 2.8 in biotin-treated microarrays were identified as candidate-positive proteins.

The Z-Score calculation formula is shown below.

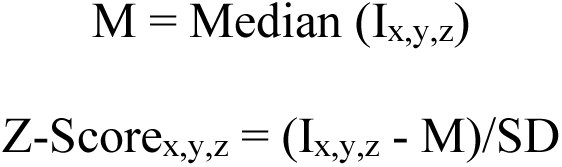

In which, M represents the median of all protein site intensities (I_x,y,z_), I_x,y,z_ denotes the raw signal intensity of each protein site on the chip. x is the column number, y is the row number, and z is the sequence number of the block on the chip. Z-Score_x,y,z_ refers to the Z-score value of each protein site within the chip, and SD represents the standard deviation of all I_x,y,z_ values.

### SPR biosensing for IgG1-Fc and nostrin binding affinity

Interactions of insulin-129 with surface-immobilised mouse IgG1-Fc (Medlife) and nostrin (Canspec China) were determined using surface plasmon resonance (SPR). Experiments were performed at 25 °C on a BIAcore 1K using CM5 sensor chips. Data were analysed using BIAcore 1K Evaluation software (Cytiva) following the manufacturer’s instructions. In brief, mouse IgG1-Fc and nostrin were immobilised to the cell surface by pre-activation. The cell was then blocked with 1 M ethanolamine. A neighbouring aisle that served as a reference was similarly activated and blocked. Both of the aisles were then equilibrated with DPBS (pH 7.4). Molecule stock solution was diluted to a series of concentrations in DPBS and was flowed at 10 μL min for 150 s in each run. At the end of each flow, cells were regenerated for 5 min with 50 mM NaOH solution at 10 μL/min. For SPR biosensing under different glucose concentrations, the running buffer and sample solution were added with different amounts of glucose before the test. Data from the sample cell were collected using Biacore Insight (version 2.0, Cytiva), and were subtracted from those from the reference cell. Association and dissociation constants were obtained by global fitting of the data to a 1:1 Langmuir binding model using BIAcore 1K Evaluation software (Cytiva, USA).

Human IgG1-Fc and nostrin were tested in the same way.

### Enzyme degradation assay

Proteolytic stability of insulin-129 and RHI (0.6 mM, 10 μL) toward cathepsin D (6.25 U/mL, 3 μL) was measured in citrate-phosphate buffer (pH 4.0) and at 37°C, with a final volume of 300 μL ^48,49^. At various time points (0, 3, 5, 15, 30, and 60 min), samples were quenched with a triple volume of acetonitrile and cooled to 0°C. Insulin-129 and RHI were analysed by the Shimadzu Prominence Ultra-Fast Liquid Chromatography System (Shimadzu Corporation, Japan), which was coupled with an AB Sciex 5500+ QTRAP mass spectrometer consisting of an ESI source (AB SCIEX, USA). The curve was fitted using one-phase decay model. Half-lives (t_1/2_) were obtained from the curves.

### Computational simulations of binding conformations

AlphaFold 3 was used to predict the interactions between target proteins (IgG1-Fc and nostrin) and insulin-129 ^50^. The process of covalent binding is typically guided by initial non-covalent interactions, where the ligand-protein first binds to the receptor protein in a favourable conformation, facilitating subsequent covalent bond formation. Thus, AlphaFold 3 was initiated to simulate the non-covalent interaction between the target proteins and insulin-129, predicting potential binding conformations and constructing a preliminary non-covalent interaction model. Specifically, the full sequences of the target proteins were retrieved from the UniProt and incorporated into the sequences section of a JSON input file, alongside the sequences of insulin’s A and B chains. The chemical structure of the phenylboronic acid group was defined in the ligand section using custom CCD mmCIF format, and its covalent bonding with the A1 and B29 residues of insulin was specified in the bondedAtomPairs section. Subsequently, based on the simulation result and the chemical property of the phenylboronic acid group, possible covalent binding sites were identified. Further modelling of these sites was conducted using AlphaFold 3, where the bondedAtomPairs field was used to define covalent interactions between the boronic acid group and the reactive center of the target protein. This process simulated the covalent interaction between the modified insulin and the target proteins, optimising the stable conformation of the resulting complexes.

### The type 1 diabetes treatment study in mice

All animal procedures were performed following the Guidelines for Care and Use of Laboratory Animals of Zhejiang University (Protocol No. ZJU20241113 and No. ZJU20240105). C57BL/6 mice (six to eight weeks) were purchased from Hangzhou Medical College. The type 1 diabetic mice were induced *via* streptozotocin (STZ) administration at a dose of 120 mg/kg. The diabetic mice were maintained with a standard diet in cycles of 12 h light and 12 h dark, allowing free access to food and water during the experiment. Diabetic mice were grouped (*n* = 5) and injected subcutaneously with insulin glargine (40 and 400 U/kg) as a commercial insulin control. All insulin analogues synthesised in this study were initially evaluated for efficacy by subcutaneous injection into diabetic mice at 4 mg/kg. Insulin-129 was subsequently tested at 8, 12, 16, 20, and 24 mg/kg to investigate the dose–efficacy relationship.. As a control, insulin glargine at a dose of 40 U/kg was injected every 24 hours for seven consecutive days respectively. Also, the blood glucose (BG) of diabetic mice without treatment was continuously monitored for one month. BG was measured using a glucose meter (Aviva, ACCU-CHEK).

### Intraperitoneal glucose tolerance test in mice

Diabetic mice treated with insulin-129 at a dose of 12 mg/kg (*n* = 5) were subjected to intraperitoneal glucose administration (1.5 g/kg) at 1 day and 5 days post-treatment. Healthy mice (*n* =5) were used as the control group. The BG was measured at different intervals. Aera under the curve (AUC) was calculated using the GraphPad Prism (version 10.0).

### Long-term therapy of insulin-129 as weekly formulation

Diabetic mice were grouped (*n* = 5) and injected subcutaneously with insulin-129 at a dose of 10 mg/kg for the first injection, followed by three injections with a dose of 4.8 mg/kg every 168 hours. The BG was measured at different intervals.

### *In vivo* glucose-responsive insulin release in mice

Diabetic mice treated with insulin-129 at a dose of 12 mg/kg (*n* = 5) were intraperitoneally injected with glucose (3.0 g/kg) at 1 day and 5 days post-treatment. Healthy mice (*n* =5) and diabetic mice treated with insulin glargine (400 U/kg, *n* = 5) were used as the control groups. The BG was measured at different intervals. Blood samples (20 μL) were collected for BG measurement and plasma extraction. The plasma insulin levels were measured using a human insulin ELISA kit (Elabscience, catalogue no. E-EL-H2665) or a mouse insulin ELISA kit (EZassay, catalogue no. MS200).

### Pharmacokinetic studies in mice

The pharmacokinetics of both transvascular and extravascular administration were investigated. For transvascular administration, healthy, non-fasted awake C57BL/6J mice (*n* = 5) were given a single intravenous injection of insulin-129 (1 nmol/kg, dissolved in PBS, pH 7.4) in the tail vein, and plasma concentration-time profiles were followed for 21 days after dosing with blood sampling at predetermined intervals. For extravascular administration, diabetic mice treated with insulin-129 (*n* = 5) were given a single subcutaneous injection of insulin-129 (12 mg/kg), and plasma concentration-time profiles were followed for 17 days after dosing with blood sampling at predetermined intervals. The plasma concentrations of insulin-129 were analysed by a human insulin ELISA kit (Elabscience, catalogue no. E-EL-H2665).

Plasma concentration−time profiles were analysed by two-compartmental pharmacokinetic analysis in WinNonlin (Certara, CA). Calculations were performed using individual concentration−time values from each animal.

### Isoglycaemic clamp studies in mice

Isoglycaemic clamp studies were conducted to establish euglycaemic and hyperglycaemic conditions in healthy mice and STZ-induced type 1 diabetic mice, respectively ^51,52^. Prior to glucose administration, both groups of vein-cannulated healthy and diabetic mice were given a recovery period of five to seven days. After a six-hour fasting period, the venous catheters were externalised and connected to a glucose infusion system. Glucose was infused either one day after administering insulin-129 at a dosage of 12 mg/kg or immediately following the injection of insulin glargine at 5 U/kg. BG levels were monitored every ten minutes using a glucose meter (Aviva, ACCU-CHEK), facilitating adjustments to the glucose infusion rate to maintain BG concentrations around 100 mg/dL for healthy mice and approximately 400 mg/dL for diabetic mice over a three-hour duration.

### The type 1 diabetes treatment study in minipigs

The Bama minipigs (six-month-old) were injected intravenously with STZ at a dose of 150 mg/kg to induce type 1 diabetes. The body weights of Pig-I, Pig-II, and Pig-III are 60.4, 54.9 and 47.6 kg, respectively. After BG was stabilised for one month, diabetic minipigs were used for treatment. The minipigs were fed twice a day, either with or without treatment. Insulin glargine was injected once a day for usual BG control. The BG was monitored by a continuous glucose monitoring system (CGMS, SIBIONICS). The administration of insulin glargine was paused for 48 hours before evaluating the BG control efficacy, and diabetic minipigs were subcutaneously injected with insulin-129 at 0.16, 0.36, and 0.63 mg/kg for Pig-I, Pig-II and Pig-III, respectively.

### Pharmacokinetic studies in minipigs

Diabetic minipigs of Pig-I, Pig-II, and Pig-III were subcutaneously injected with insulin-129 at 0.16, 0.22, and 0.32 mg/kg, respectively. Blood (2 mL) was drawn at specified time points using serum collection tubes (BD Vacutainer) for serum separation and collection. The serum insulin levels were measured using a human insulin ELISA kit ((Elabscience, catalogue no. E-EL-H2665)). The individual serum insulin concentration−time profile was analysed using a two-compartmental pharmacokinetic analysis in WinNonlin (Certara, CA).

### *In vivo* blood glucose-responsive insulin release in minipigs

At one day posttreatment, dextrose (10 wt %) solution was infused at a rate of 1 litre/hour until the BG of the minipigs was above 300 mg/dL, while the minipigs were kept anaesthetised using isoflurane. Blood was collected from the external jugular vein before and during the experiment at scheduled intervals. Blood (2 mL) was placed in serum collecting tubes (BD Vacutainer). Serum was collected following the protocol. Serum insulin levels were measured by a human insulin ELISA kit (Elabscience, catalogue no. E-EL-H2665).

### Application of insulin-129 in a dual-closed-loop fully automated insulin delivery system

The dual-closed-loop fully automated insulin delivery system included a CGMS, Android APS equipped with the algorithm, Danner R insulin pump (Sooil, Korea), and insulin-129 (3 mg/mL, in PBS at pH 7.4). The main parameters of the algorithm module involved administration interval, dose, and BG at insulin infusion initiation. When two consecutive BG points obtained by the CGMS were higher than 150 mg/dL and the interval since the last dose was longer than 2 hours, the insulin pump delivered a bolus of insulin-129 at 0.2, 0.3 or 0.2 mg for Pig IV, V and VI. No insulin-129 was delivered under all other conditions. CGMS recorded BG every five minutes and uploaded it to the host device (Pixel 4 XL, Google) installed with Android APS. The pump received the command and determined the infusion of insulin-129.

In the insulin lispro-applied single-closed-loop fully automated insulin delivery system, the algorithm of Android APS was set the same as the insulin-129 treated group, except that the bolus doses of insulin lispro were set as 0.24, 0.14 and 0.12 mg for Pig VII, VIII and IX. The start-up was the same as insulin-129 treatment group.

### Biodistribution of insulin-129 in mice

Cy5-labelled insulin-129 was administered to diabetic mice at a dose of 10 mg/kg. At designed intervals, the mice were euthanised and the brain, heart, lung, liver, spleen, kidney, bladder, fat, and muscle were dissected for *ex vivo* fluorescence imaging. Diabetic mice receiving Cy5-labelled RHI and free Cy5 were taken as control groups. The Cy5-labelled RHI was quantified using a microplate reader (Synergy H1, BioTek, USA) to ensure the same injection fluorescence intensity as Cy5-labelled insulin-129. The free Cy5 was injected according to the amount of Cy5 contained in Cy5-labelled insulin-129.

### Intra-tissue distribution of insulin-129 in mice

Cy5-labelled insulin-129 (10 mg/kg) was administered to diabetic mice. Three days post-injection, the mice were euthanised and major organs were harvested and prepared as cryosections. The sections were fixed with cold acetone and subsequently subjected to immunofluorescence staining. Immunofluorescence staining of CD31 (ABclonal, catalogue no. A23701, 1:100) was performed following standard protocol. The images were collected using a digital slice scanner VS200 (Olympus, Japan). Olympus Image Viewer software (version 4.1) was used for analysis.

### Toxicity evaluation in mice

Diabetic mice were injected subcutaneously with insulin-129 or PBS (*n* = 5). For a single injection of insulin-129 at a dose of 12 mg/kg and PBS, the whole blood and serum were collected seven days post-treatment from the treated mice. For insulin-129 as a weekly formulation, we collected whole blood and serum at 28 days after multiple injections. Whole blood and serum were also collected from healthy mice as a control. Blood and serum samples were analysed for routine tests, including blood cell counting and measurements of aspartate transaminase (AST), alanine transaminase (ALT), alkaline phosphatase (ALP), albumin (ALB), blood urea nitrogen (BUN), and creatinine levels.

### Histomorphological observation of major tissues in mice

Insulin-129 (10 mg/kg) was administered to diabetic mice. One week after the injection, the mice were euthanised and their major organs were collected. The tissues were fixed in 4% PFA, embedded in paraffin, and sectioned prior to hematoxylin and eosin (H&E) staining. Digital images were captured using a VS200 digital slice scanner (Olympus), and the analysis was performed with Olympus Image Viewer software (version 4.1).

### Statistics and reproducibility

Statistical significance in all figures was calculated by two-tailed unpaired Student’s *t*-test. \**P* < 0.05, \*\**P* < 0.01, \*\*\**P* < 0.001, \*\*\*\**P* < 0.0001, ns, not significant. All *P* value analyses were performed on GraphPad Prism (version 10.0). A minimum of three biological replicates were analysed for quantitative analysis. All data were means ± SD.

## Data availability

Source data are provided in this paper. All data supporting this study’s findings are presented in the paper and the supplementary information.

