## Supplemental figures for "An ultra-long acting insulin enables glucose-synchronised release"

**Contents**

**Supplementary Fig. 1 |** Characterization of insulin-129, insulin-1, and insulin-29.

**Supplementary Fig. 2 |** Preliminary pharmacodynamics and mechanism evaluation of insulin-129.

**Supplementary Fig. 3 |** The docking analysis of insulin-129 with IgG1-Fc and nostrin.

**Supplementary Fig. 4 |** *In vivo* treatment of insulin-129 in mice.

**Supplementary Fig. 5 |** Efficacy assessment of insulin-129 in minipigs.

**Supplementary Fig. 6 |** Biodistribution of free Cy5 and Cy5-labelled RHI and biosafety of insulin-129.

**Supplementary Fig. 7 |** Interactions of insulin-129 with human IgG1-Fc and nostrin.

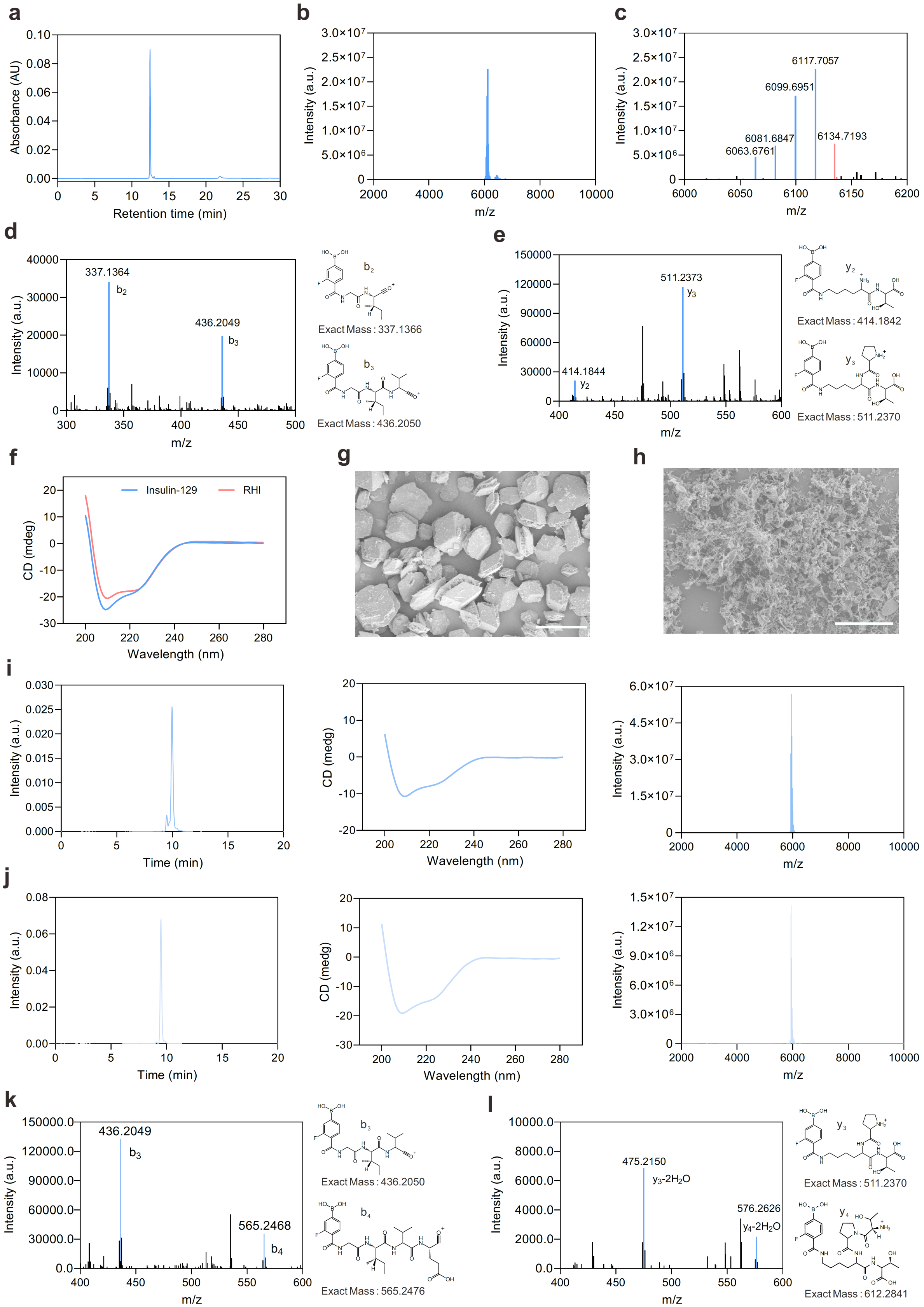
a
Absorbance (AU)
Retention time (min)
b
Intensity (a.u.)
m/z
c
6117.7057
6099.6951
6081.6847
6063.6761
6134.7193
d
337.1364
b2
436.2049
b3
Exact Mass : 337.1366
Exact Mass : 436.2050
e
511.2373
y3
414.1844
y2
Exact Mass : 414.1842
Exact Mass : 511.2370
f
Insulin-129
RHI
CD (mdeg)
Wavelength (nm)
g
h
i
Intensity (a.u.)
Time (min)
CD (medg)
Wavelength (nm)
m/z
j
Intensity (a.u.)
Time (min)
CD (medg)
Wavelength (nm)
m/z
k
436.2049
b3
565.2468
b4
Exact Mass : 436.2050
Exact Mass : 565.2476
l
475.2150
y3-2H2O
576.2626
y4-2H2O
Exact Mass : 511.2370
Exact Mass : 612.2841

**Supplementary Fig. 1 | Characterization of insulin-129, insulin-1, and insulin-29. a,** Reversed-phase high-performance liquid chromatography (RP-HPLC) of insulin-129 (absorbance at 280 nm). **b,** High-resolution mass spectrometry (HRMS) of insulin-129. **c,** Local magnification of (**b**) in the m/z range of 6000 to 6200. The labelled peaks represented the molecule ion peak (red) and its four dehydration peaks (blue). From right to left, 1 to 4 $H_2O$ were dropped, respectively. **d,e,** Tandem mass spectrometry (MS/MS) spectra of A Chain (**d**) and B Chain (**e**) for insulin-129. The presence of diagnostic b and y ions in the MS/MS spectra confirmed modification sites of 4-carboxy-3-fluorophenylboramide at A1 and B29 residues. **f,** Circular dichroism (CD) spectra of insulin-129 and recombinant human insulin (RHI) at a concentration of 0.2 mg/mL. **g,** Representative scanning electron microscope (SEM) image of RHI. Scale bar, 50 μm. **h,** Representative SEM image of insulin-129. Scale bar, 25 μm. **i,** RP-HPLC chromatogram, CD spectra, and HRMS of insulin-1, respectively (from left to right). **j,** RP-HPLC chromatogram, CD spectra, and HRMS of insulin-29, respectively (from left to right). **k,** MS/MS spectrum of A Chain for insulin-1. The presence of diagnostic b and y ions in the MS/MS spectrums confirmed the exact modification sites 4-carboxy-3-fluorophenylboramide are at the A1 position. **l,** MS/MS spectrums of B Chain for insulin-29. The presence of diagnostic b and y ions in the MS/MS spectrums confirmed the exact modification sites 4-carboxy-3-fluorophenylboramide are at the B29 position.

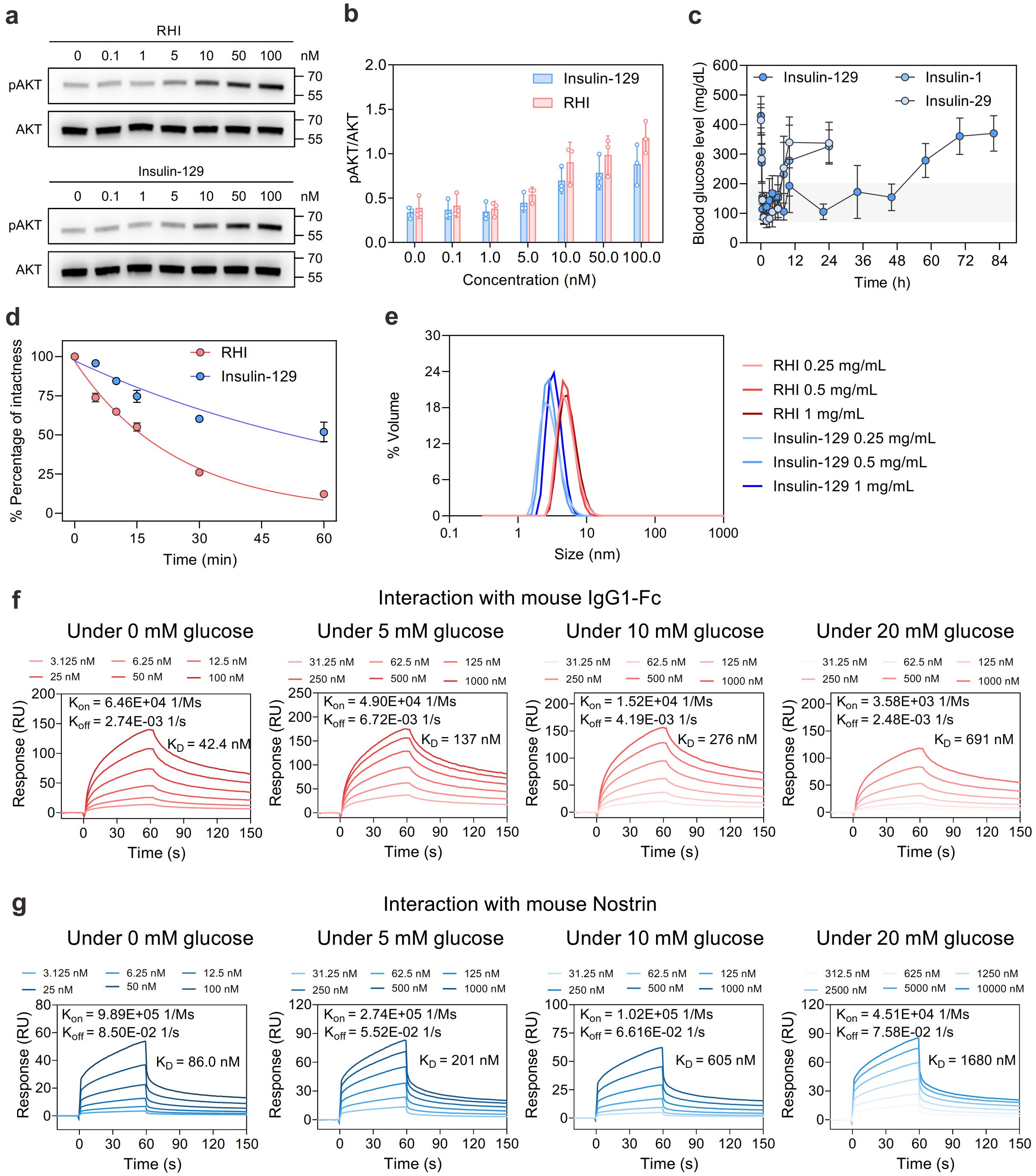


**Supplementary Fig. 2 | Preliminary pharmacodynamics and mechanism evaluation of insulin-129. a,** Phosphorylation of AKT signalling by the indicated concentrations of RHI or insulin-129 for 10 min in HepG2 cells. **b,** The ratio of pAKT to AKT in HepG2 cells stimulated by RHI or insulin-129. Data points are means ± s.d. ($n = 3$) and individual points. **c,** Blood glucose (BG) level of diabetic mice treated with subcutaneously injected insulin-1, insulin-29, and insulin-129 (4 mg/kg). Data points are mean ± s.d. ($n = 5$). **d,** The percentage of intact insulin-129 and RHI against time were plotted using GraphPad Prism (version 10). The curve was fitted using a one-phase decay model. Data points are means ± s.d. ($n$ = 3) **e,** The dynamic light scattering measurement by volume was performed at 25 °C in 1× PBS buffer with concentrations of 0.25, 0.5, and 1 mg/mL. **f,** The surface plasmon resonance (SPR) results of insulin-129 with IgG1-Fc under different glucose concentrations of 0, 5, 10 and 20 mM. **g,** The SPR results of insulin-129 with nostrin under different glucose concentrations of 0, 5, 10 and 20 mM.

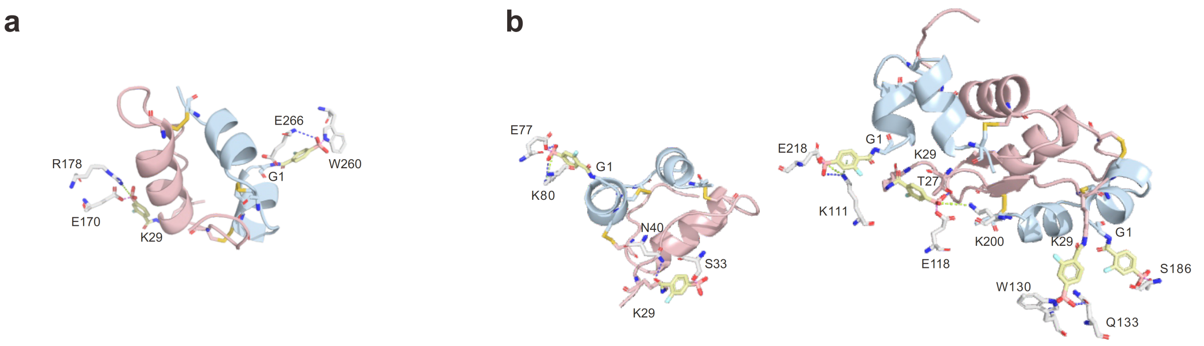


**Supplementary Fig. 3 | The docking analysis of insulin-129 with IgG1-Fc and nostrin. a,** Simulated binding conformation of insulin-129 with IgG1-Fc. FPBA modified at G1 and K29 could form reversible covalent interactions with the W260 and E170 residues of IgG-Fc, respectively. **b,** Simulated binding conformations of insulin-129 with nostrin. The FPBA moieties modified at residues G1 and K29 on insulin-129 could form reversible covalent interactions with distinct nostrin residues, specifically E77 and S33 (left), E218 and E118 (middle), and W130 and S186 (right), respectively. The covalent interactions are depicted as violet dashed lines, salt bridges are represented by lime dashed lines, hydrogen bonds are indicated with blue dashed lines and cation-π interactions are shown as smudge dashed lines.

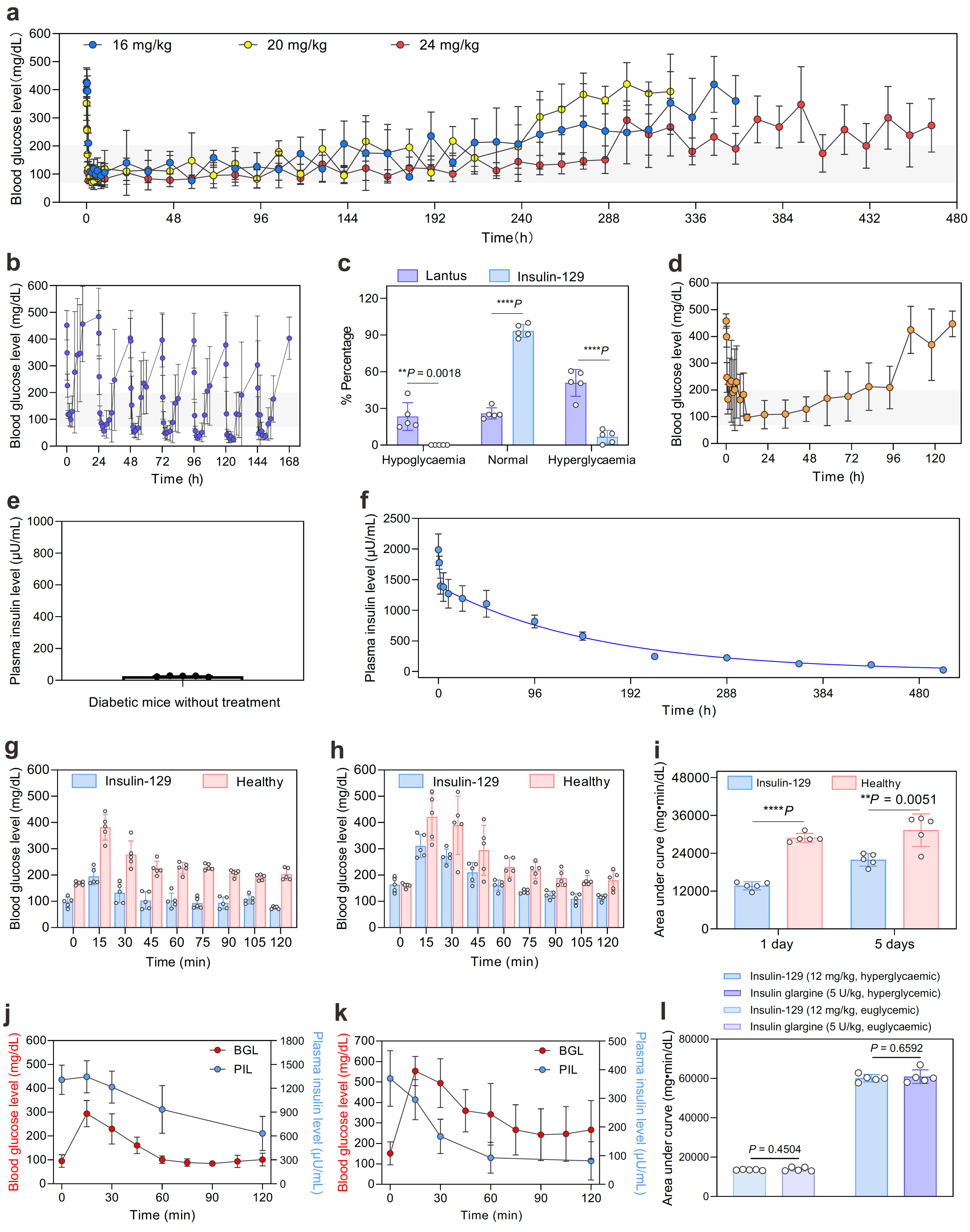


**Supplementary Fig. 4 | *In vivo* treatment of insulin-129 in mice. a,** BG of diabetic mice treated with subcutaneously injected insulin-129 (16, 20, and 24 mg/kg). Data points are means ± s.d. ($n = 5$). **b,** BG of diabetic mice treated with seven injections of Lantus (40 U/kg, one injection per day). Data points are means ± s.d. ($n$ = 5). **c,** Hypoglycaemia, normal, and hyperglycaemia are defined as BG < 70 mg/dL, BG between 70 and 200 mg/dL, and BG > 200 mg/dL, respectively. Data points are means ± s.d. ($n$ = 5) and individual points. Statistical time for the insulin-129-treated group (12 mg/kg) and Lantus-treated group were 0-178 h and 0-168 h, respectively. Unpaired two-tailed Student's $t$-test was used for statistical analysis. **$P < 0.01$, ****$P < 0.0001$. **d,** BG of diabetic mice treated with subcutaneously injected insulin icodec (400 U/kg). Data points are means ± s.d. ($n$ = 5). **e,** Plasma insulin-129 levels of diabetic mice without treatment. Data are shown as mean ± s.d. ($n$ = 5) and individual points. **f,** Pharmacokinetics of insulin-129 after a single intravenous injection in healthy mice (1 nmol/kg). Data points are mean ± s.d. ($n$ = 5). The curve was fitted to the two-compartment model. **g,h,** Glucose was intraperitoneally injected (1.5 g/kg) to diabetic mice received insulin-129 injection for one (**g**) and five days (**h**). Data points are means ± s.d. ($n$ = 5) and individual points. **i,** Area under curve statistics for (**g**) and (**h**). Data points are means ± s.d. ($n$ = 5) and individual points. Unpaired two-tailed Student's $t$-test was used for statistical analysis. **$P < 0.01$, ****$P$ <0.0001. **j,k,** Glucose was intraperitoneally injected (3.0 g/kg) after diabetic mice received insulin glargine (400 U/kg) for three hours (**j**) and six hours (**k**). The BG and insulin glargine levels were tested at different intervals. Data points are means ± s.d. ($n$ = 5). **l,** Area under curve statistics for BG-time curves of all groups in isoglycaemic clamp studies. Data points are means ± s.d. ($n$ = 5) and individual points. Unparied two-tailed Student's $t$-test was used for statistical analysis.

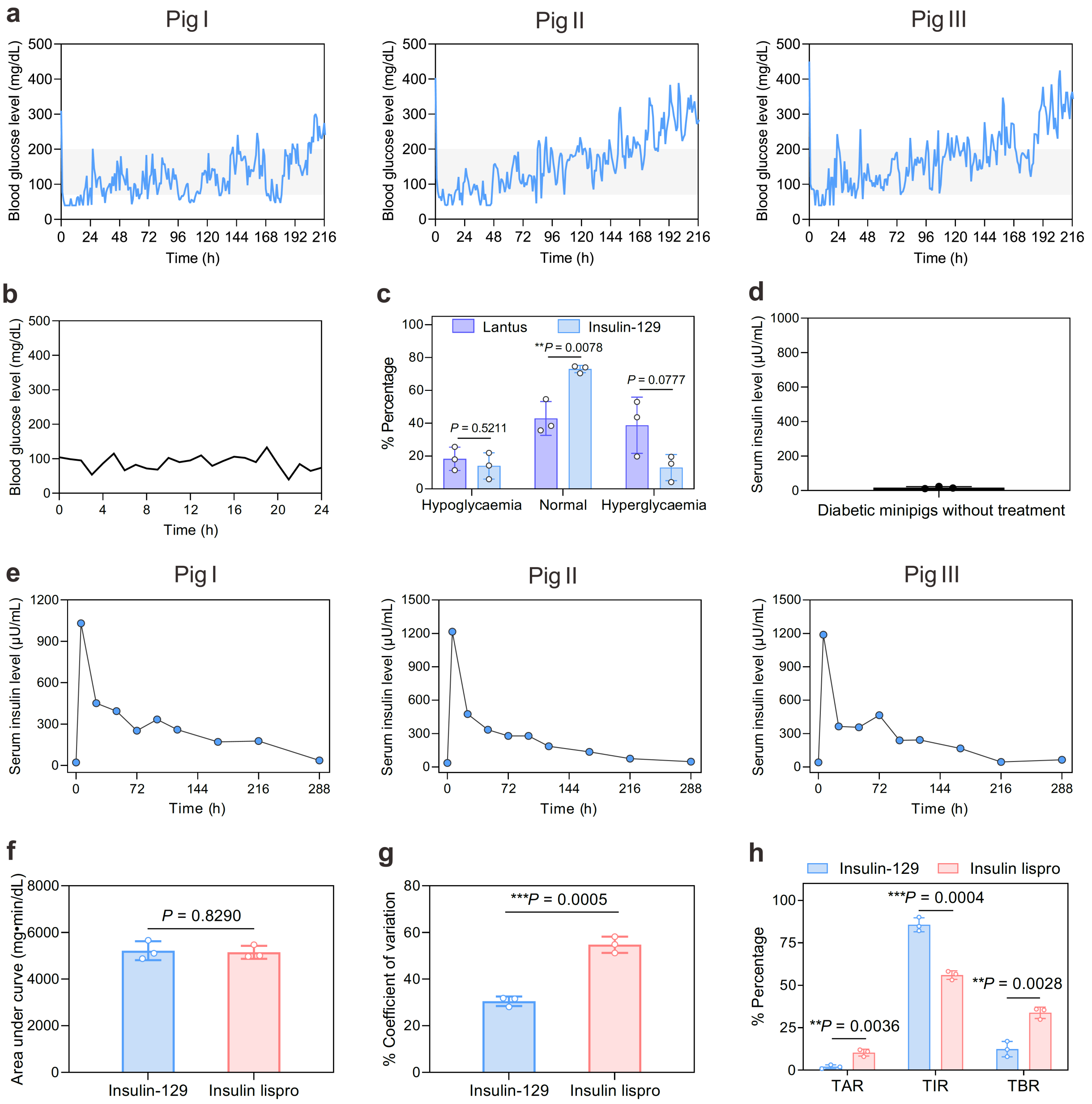


**Supplementary Fig. 5 | Efficacy assessment of insulin-129 in minipigs. a,** BG of type 1 diabetic minipigs treated with one injection of insulin-129 (0.16, 0.36 and 0.63 mg/kg for Pig I, Pig II and Pig III, respectively). **b,** Representative daily BG profile of a healthy pig. **c,** Hypoglycaemia, normal, and hyperglycaemia are defined as BG < 70 mg/dL, BG between 70 and 200 mg/dL, and BG > 200 mg/dL, respectively. Data points are means ± s.d. (*n* = 3) and individual points. Statistical time for the insulin-129-treated group and Lantus-treated group were 0-168 h and 0-24 h, respectively. Unpaired two-tailed Student's *t*-test was used for statistical analysis. $^{**}P < 0.01$. **d,** Serum insulin-129 levels of diabetic minipigs without treatment. Data are shown as mean ± s.d. (*n* = 3) and individual points. **e,** The minipigs were treated with subcutaneously injected insulin-129 at a dose of 0.16, 0.22 and 0.32 mg/kg to Pig I, Pig II and Pig III, respectively. **f,** Area under curve statistics for BG-time curves of insulin-129-treated and insulin lispro-treated groups. Data points are means ± s.d. ($n = 3$) and individual points. A two-tailed Student's $t$-test was used for statistical analysis. **g,** Coefficient of glucose variation. Data points are means ± s.d. ($n = 3$) and individual points. A two-tailed Student's $t$-test was used for statistical analysis. A two-tailed Student's $t$-test was used for statistical analysis. ****$P < 0.001$. **h,** Time in range (70-180 mg/dL, TIR), time above range (>180 mg/dL, TAR) and time below range (<70 mg/dL, TBR). Data points are means ± s.d. ($n = 3$) and individual points. A two-tailed Student's $t$-test was used for statistical analysis. **$P < 0.01$, ***$P < 0.001$.

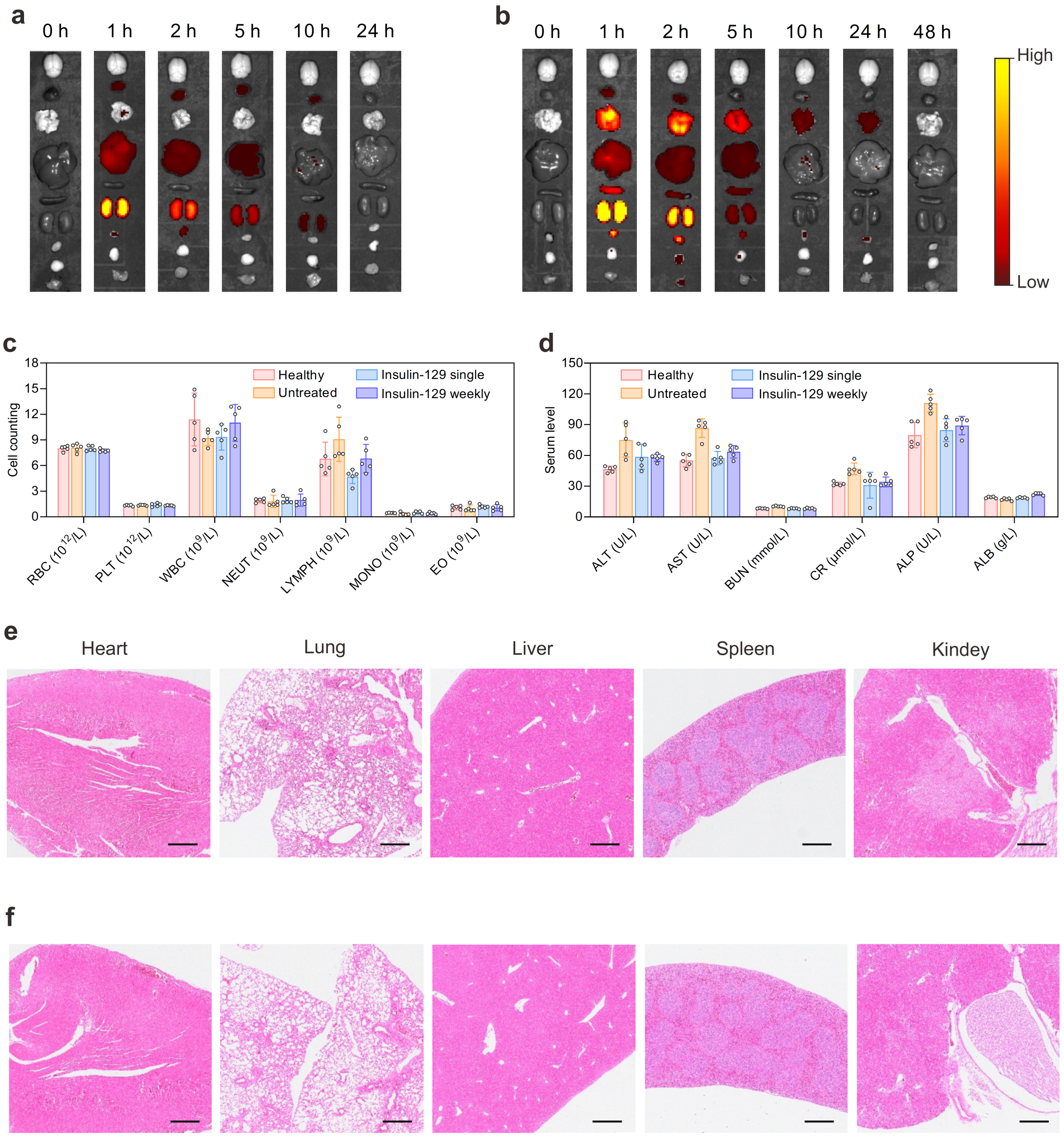


**Supplementary Fig. 6 | Biodistribution of free Cy5 and Cy5-labelled RHI and biosafety of insulin-129. a,** Representative *ex vivo* fluorescence imaging of type 1 diabetic mice after subcutaneous injection of free Cy5. **b,** Representative *ex vivo* fluorescence imaging of type 1 diabetic mice after subcutaneous injection of Cy5-labelled RHI. The major organs or tissues (from top to bottom) included the brain, heart, lung, liver, spleen, kidney, bladder, fat and skeleton muscle. The color scale indicates the radiant efficiency from $1.00 \times 10^8$ to $3.00 \times 10^9$ photons per s per $cm^2$ per sr. The Cy5-labelled RHI was quantified using a microplate reader to ensure the same injection fluorescence intensity as Cy5-labelled insulin-129. The free

Cy5 was injected according to the amount of Cy5 contained in Cy5-labelled insulin-129. **c,d,** The blood cell count (**c**) and serum level of major biochemical parameters (**d**) in diabetic mice with subcutaneously-injected insulin-129. Diabetic mice receiving PBS and healthy mice were used as control groups. Whole blood was obtained one week after a single injection of insulin-129. Data points are means ± s.d. ($n$ = 5) and individual points. RBC, red blood cell; PLT, platelet; WBC, white blood cell; NEUT, neutrophil; LYMPH, lymphocyte; MONO, monocyte; EO, eosinophil. ALT, alanine transaminase; AST, aspartate transaminase; BUN, blood urea nitrogen; CR, creatinine; ALP, alkaline phosphatase; ALB, albumin. **e,** Representative H&E staining of major organs at 7 days after the subcutaneous administration of insulin-129 at a dose of 10 mg/kg. **f,** Representative H&E staining of major organs at 7 days after the subcutaneous administration of PBS (pH 7.4). Scale bar, 500 μm.

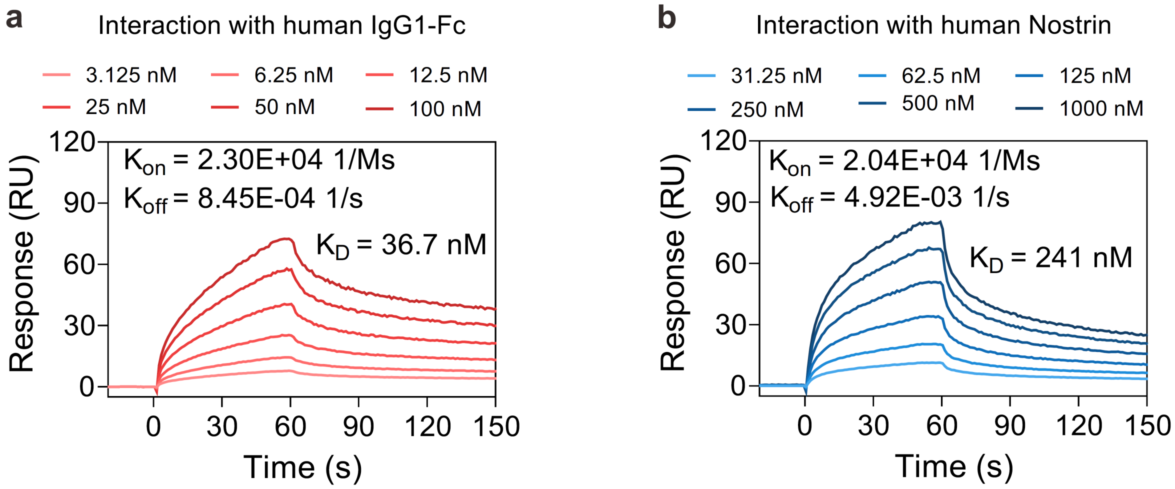


**Supplementary Fig. 7 | Interactions of insulin-129 with human IgG1-Fc and nostrin. a,** The SPR results of insulin-129 with human IgG1-Fc. **b,** The SPR results of insulin-129 with human nostrin.
